# The cell wall polymer initiates plasma membrane partitioning in mycobacteria

**DOI:** 10.1101/2022.06.12.495848

**Authors:** Takehiro Kado, Zarina Akbary, Daisuke Motooka, Ian L. Sparks, Emily S. Melzer, Shota Nakamura, Enrique R. Rojas, Yasu S. Morita, M. Sloan Siegrist

## Abstract

Lateral partitioning of proteins and lipids shapes membrane function. In model membranes, partitioning can be influenced by interactions with other membranes and solid supports. While cellular membranes can departition in response to various perturbations, including disruption of bilayer-extrinsic structures, the mechanisms by which they partition *de novo* are largely unknown. The plasma membrane of *Mycobacterium smegmatis* can be spatially and biochemically departitioned by the fluidizing agent benzyl alcohol. By screening for mutants that are sensitive to benzyl alcohol, we show that the bifunctional cell wall synthase PonA2 promotes membrane partitioning and cell growth upon fluidizer washout. PonA2’s role in membrane repartitioning and regrowth depends solely on its conserved transglycosylase domain. We find that the cell wall polymer, but not active cell wall polymerization, is critical for membrane partitioning. Our work highlights a key initiating role for bilayer-extrinsic structures in patterning cellular membranes.

## Introduction

Biological membranes are heterogeneous mixtures of lipids and proteins (Bernardino de la Serna et al., 2016; Singer and Nicolson, 1972). In eukaryotic cells, membrane domains host functions such as signal transduction, membrane sorting, protein processing, and virus trafficking (Goñi, 2019; Simons and Sampaio, 2011). Bacterial membrane domains have long been suggested (Epand and Epand, 2009; Matsumoto et al., 2006), but the molecular mechanisms and physiological significance of membrane compartmentalization in these organisms are only beginning to emerge. To date, the best-studied example is functional membrane microdomains (FMM), which are present in many bacterial species and are, like their eukaryotic counterparts, more liquid-ordered, or rigid, than the surrounding plasma membrane (López and Kolter, 2010). FMMs contribute to diverse biological functions including signaling, cell morphology maintenance, and biofilm formation (Bach and Bramkamp, 2013; Dempwolff et al., 2012a, 2012b; Mielich-Süss et al., 2013; Yepes et al., 2012; Zielińska et al., 2020). In contrast to FMMs, regions of increased fluidity (RIF) are more liquid-disordered than the surrounding membrane . RIFs are also present in many bacterial species but their function(s) remain relatively unexplored (Gohrbandt et al., 2022; Molohon et al., 2016; Strahl et al., 2014; Wenzel et al., 2018).

We have shown that bacteria of the pole-growing genus *Mycobacterium—*which includes both pathogens, such as *M. tuberculosis,* and saprophytes, such as *M. smegmatis*—partition their plasma membranes into the intracellular membrane domain (IMD) and conventional plasma membrane, which is termed the PM-CW as it is tightly associated with the cell wall (García-Heredia et al., 2021; Hayashi et al., 2016, 2018; Morita et al., 2005; Puffal et al., 2022). The biophysical nature of the IMD is not known but may be more liquid-disordered than the surrounding PM-CW given that the IMD is enriched in enzymes that act on membrane-fluidizing polyprenols and lipid intermediates carrying polymethylated fatty acids (García-Heredia et al., 2021; Hayashi et al., 2016, 2018; Janas et al., 1994; Morita et al., 2005; Puffal et al., 2022; Schroeder et al., 1987; Valtersson et al., 1985; Vigo et al., 1984; Wang et al., 2008). Biosynthetic pathways that are partitioned across the IMD and PM-CW include both the well-conserved, *e.g*., cell wall peptidoglycan and plasma membrane phosphatidylethanolamine (PE), and the mycobacteria-specific, *e.g*., plasma membrane glycolipids phosphatidylinositol mannosides (PIMs) and outer membrane phthiocerol dimycocerosate (PDIM) (García-Heredia et al., 2021; Hayashi et al., 2018; Morita et al., 2005; Puffal et al., 2018, 2022). While the functional role of partitioning has not yet been demonstrated for most of these pathways, perturbations that departition the membrane also dampen peptidoglycan precursor production (Garcia-Heredia 2021), indicating that lateral partitioning may promote membrane-bound enzymatic reactions.

Partitioning in model membranes can be controlled by both bilayer-intrinsic factors, *i.e.,* protein (Yuan et al., 2021) a lipid composition (Beales et al., 2005; Korlach et al., 1999; Veatch and Keller, 2003), as well as bilayer-extrinsic factors, including temperature and interactions with other membranes and/or solid supports (Gordon et al., 2008; Subramaniam et al., 2013). It has been experimentally challenging to perform analogous experiments in the more-complex membranes of living cells (Gohrbandt et al., 2022). Therefore, most work on cellular membrane partitioning has concentrated on defining maintenance factors. One factor that maintains the behavior of membrane components across different domains of life is physical connection to extracellular surfaces. For example, enzymatic removal of neuronal extracellular matrix (Frischknecht et al., 2009) or the plant or bacterial cell wall (Daněk et al., 2020; Feraru et al., 2011; Martinière et al., 2012; McKenna et al., 2019; Wagner et al., 2020) can alter some of which are normally enriched in domains. Genetic or pharmacological inhibition of cell wall synthesis sometimes, but not always, disrupts membrane partitioning or proxies thereof (Feraru et al., 2011)(Daněk et al., 2020; Feraru et al., 2011; García-Heredia et al., 2021; Hayashi et al., 2018; Wagner et al., 2020). The molecular mechanisms by which physical, bilayer-extrinsic interactions control membrane partitioning are not known.

While studying membrane departitioning can illuminate requirements for maintenance, tracking repartitioning can shed light on the requirements for *de novo* establishment. Here we develop a benzyl alcohol-induced membrane departitioning/repartitioning model to screen for factors that control partitioning in cells. We identify *ponA2* as a gene that enables *M. smegmatis* to recover from benzyl alcohol. PonA2 is a bifunctional cell wall synthase that is not required for *M. smegmatis* or *M. tuberculosis* growth but counteracts various stresses, including those known to impact membrane phase behavior, *e.g.,* heat, and others indicative of potential membrane permeability defects (Kieser et al., 2015a; Li et al., 2022; Patru and Pavelka, 2010; Vandal et al., 2008, 2009a). Post-benzyl alcohol, PonA2 stimulates *M. smegmatis* regrowth and membrane repartitioning. Unlike its roles in localizing cell wall synthesis and maintaining the cell permeability barrier, which depend on conserved transglycosylase and transpeptidase domains, or its role in maintaining cell morphology, which does not depend on either domain, PonA2’s contribution to membrane repartitioning depends solely on its conserved transglycosylase domain. Inhibition of active cell wall synthesis after fluidizer washout did not impair membrane repartitioning of wild-type *M. smegmatis*. By contrast, disruption of the cell wall glycan backbone by limited glycoside hydrolase treatment rapidly departitioned the membrane. Taken together, our data indicate that the cell wall polymer, but not cell wall polymerization, is required for efficient initiation of membrane partitioning. Our work indicates that structures outside of the bilayer can regulate *de novo* partitioning of cellular membranes.

## Results

### Reversible departitioning and repartitioning of the mycobacterial plasma membrane

We previously demonstrated that the known plasma membrane fluidizer benzyl alcohol (Friedlander et al., 1987; Ingram, 1976; Konopásek et al., 2000; Nagy et al., 2007; Strahl et al., 2014; Zielińska et al., 2020) disrupts the spatial and biochemical partitioning of the *M. smegmatis* plasma membrane and halts polar growth of the organism (García-Heredia et al., 2021). The effects of benzyl alcohol reverse within 30 minutes after removing the chemical from the growth medium (García-Heredia et al., 2021). We wondered whether the ability of *M. smegmatis* to repartition its membrane post-benzyl alcohol extended to recovery from other membrane stressors. Accordingly, we treated *M. smegmatis* expressing functional fluorescent protein fusions to IMD-associated proteins (García-Heredia et al., 2021; Hayashi et al., 2016) with four different chemicals that have membrane-disrupting activity: benzyl alcohol (Maula et al., 2009; Pope et al., 1986), dibucaine (Kinoshita et al., 2019), SDS (Kopanchuk and Rinken, 2001), and oleic acid (Kurniawan et al., 2017). IMD-associated proteins such as GlfT2, MurG and Ppm1 normally localize adjacent the sites of polar growth in mycobacteria (García-Heredia et al., 2021; Hayashi et al., 2016). Upon treatment with membrane-perturbing chemicals, we found that GlfT2 and MurG were delocalized from the subpolar region by some of the chemicals whereas Ppm1 was delocalized by all (Fig. 1A and C). After 12 hours of recovery in the absence of the chemicals, the IMD marker proteins relocalized to their subpolar positions (Fig.1B). These data suggest that *M. smegmatis* can repartition its membrane after disruption by different chemicals.

**Figure 1.**
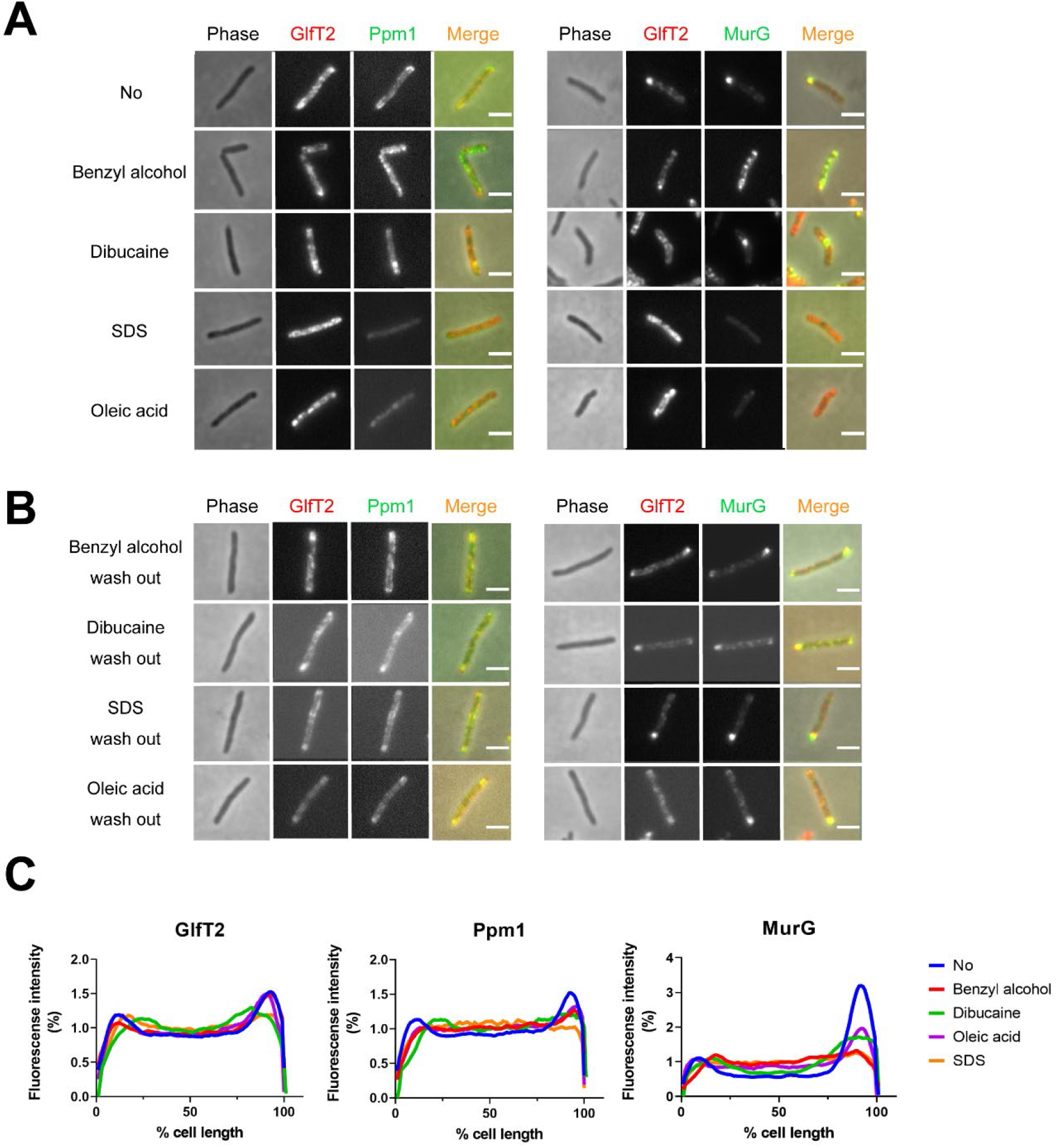
Delocalization of IMD proteins by membrane-disrupting chemicals in *M. smegmatis*. mCherry-GlfT2, Ppm1-mNeonGreem, and MurG-Dendra2 are functional fluorescent protein fusions to well-established, IMD-associated proteins (Hayashi et al., 2016, 2018; García-Heredia et al., 2021). IMD fusions were imaged after 40-minute sodium dodecyl sulfate (SDS), 40-minute oleic acid, 1-hour benzyl alcohol, or 1-hour dibucaine treatment, (A), and again 12 hours after washout, (B). Scale bars, 2.5 μm. (C) Fluorescence distributions of the fusion proteins were calculated from three independent experiments. Lines show the average of total cells (50 < n < 75). Signal was normalized to cell length and to total fluorescence intensity. Cells were oriented such that the brighter poles are on the right-hand side of the graph. See Materials and Methods for details.

### Identification of genes that promote tolerance to and/or recovery from membrane fluidization

Subpolar IMD localization correlates closely with polar cell growth (Hayashi et al., 2018). We hypothesized that the ability of *M. smegmatis* to recover from benzyl alcohol depends at least in part on its ability to reform the IMD, and therefore, genes that promote membrane partitioning would constitute a subset of the genes that enable *M. smegmatis* to tolerate and/or recover from benzyl alcohol. We performed Tn-seq (Gawronski et al., 2009; Goodman et al., 2009; Langridge et al., 2009; van Opijnen et al., 2009) to identify transposon insertions that are underrepresented 16-24 hours post-benzyl alcohol exposure compared to DMSO vehicle control. Using the TRANSIT platform (DeJesus et al., 2015), we identified six gene in which insertions that were significantly underrepresented (Fig. 2 and Table 1), suggesting that the genes might promote tolerance to and/or recovery from membrane disruption. Of the six genes, *ponA2* showed the greatest statistical significance.

**Figure 2.**
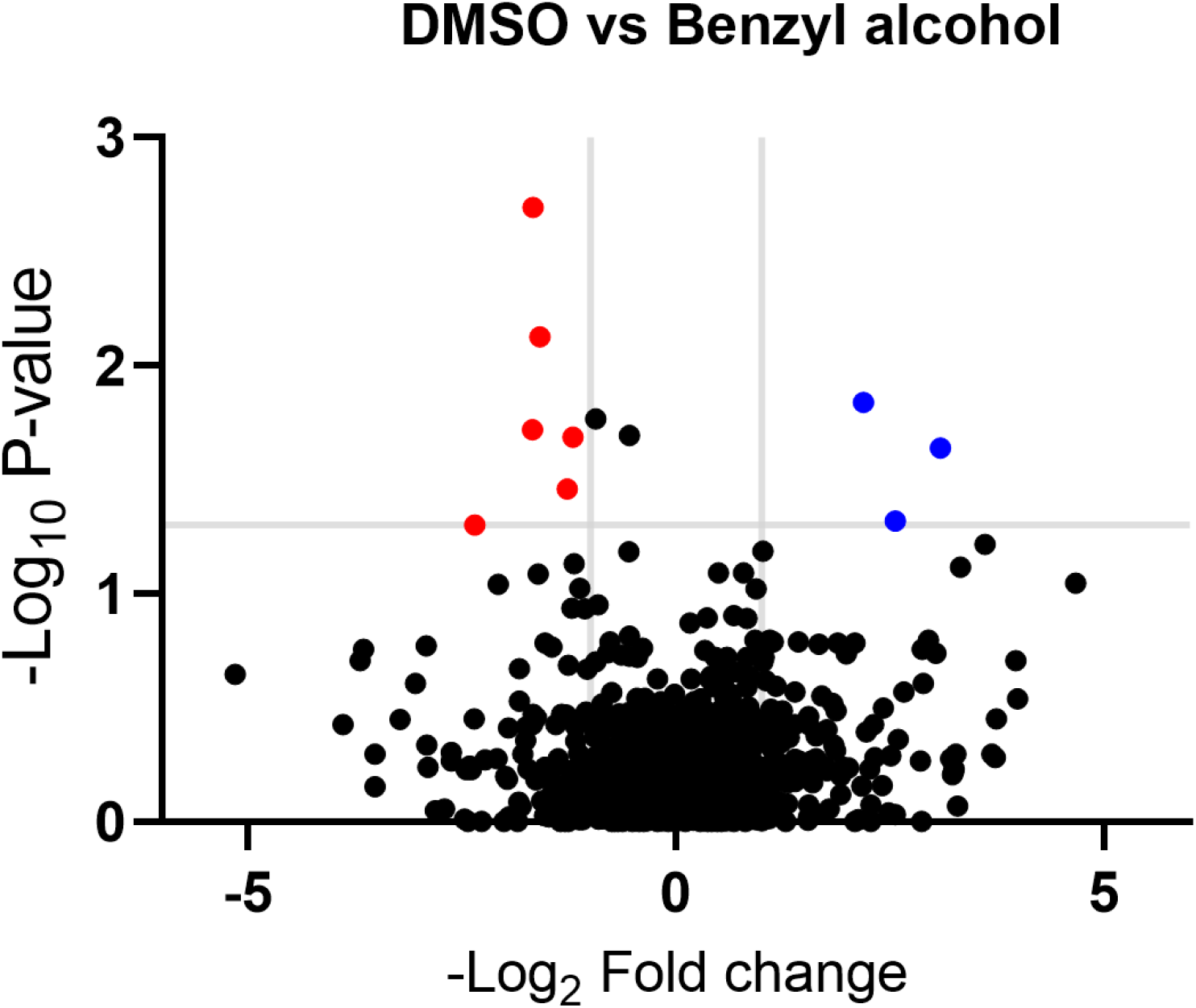
Genes in which transposon insertions are under (red, left) or over (blue, right) represented in *M. smegmatis* exposed to benzyl alcohol relative to DMSO vehicle control. A transposon library was treated with benzyl alcohol or DMSO for 1 hour. Benzyl alcohol was washed away and bacteria were resuspended in Middlebrook 7H9 growth medium. The OD_600_ was then adjusted to 0.01 and bacteria were incubated for an additional 16 -24 hours to an OD_600_ of ∼ 1.0. The library was then collected for DNA sequencing. Transposon insertion counts presented relative to counts ratio (treated/control) per gene and the corresponding P-values calculated by Mann Whitney U-test (y-axis) from n = 3 independent experiments. The horizontal grey line indicates p < 0.05; the vertical grey lines indicate 2-fold change.

**Table 1.** Genes identified by Tn-seq.

| Underrepresented in benzyl alcohol-treated *M. smegmatis* (candidates for promoting survival) |  |  |
| --- | --- | --- |
| Gene locus | Gene name | Gene description |
| MSMEG_0846c | - | putative monovalent cation/H<sup>+</sup> antiporter subunit D |
| MSMEG_2775 | *nhaA* | Na<sup>+</sup>/H<sup>+</sup> antiporter NhaA |
| MSMEG_4533c | - | sulfate-binding protein |
| MSMEG_5488c | - | DNA-binding response regulator |
| MSMEG_5781c | *pstC* | phosphate ABC transporter, permease protein PstC |
| MSMEG_6201 | *ponA2* | bifunctional transglycosylase/transpeptidase |
| Overrepresented in benzyl alcohol-treated *M. smegmatis* (candidates for impairing survival) |  |  |
| MSMEG_2768 | - | OB-fold nucleic acid binding domain-containing protein |
| MSMEG_2772 | - | amino acid permease |
| MSMEG_5694 | - | hypothetical protein MSMEG_5694 |

PonA2 is one of three bifunctional transglycosylase/transpeptidase enzymes, also known as class A penicillin-binding proteins (aPBPs), involved in cell wall peptidoglycan biosynthesis in *M. smegmatis.* We have shown that PonA1 and likely PonA2 are enriched in the PM-CW (García-Heredia et al., 2021; Hayashi et al., 2016). While PonA1 is essential for growth or viability, PonA2 and PonA3 are not (Kieser et al., 2015a; Patru and Pavelka, 2010; Vandal et al., 2008). PonA3 is not present in *M. tuberculosis* and not expressed in *M. smegmatis* under normal growth conditions (Patru and Pavelka, 2010). In contrast, PonA2 is conserved in both species, and promotes survival of *M. smegmatis* under stress conditions such as starvation or oxygen depletion; *M. tuberculosis* tolerance to heat, some antibiotics, acid, reactive oxygen and nitrogen; *M. tuberculosis* survival in some mouse backgrounds (DeJesus et al., 2017a; Kieser et al., 2015a; Li et al., 2022; Patru and Pavelka, 2010; Smith et al., 2022; Vandal et al., 2008, 2009a). As enzymatic removal of the *M. smegmatis* cell wall departitions the membrane (García-Heredia et al., 2021), we reasoned that PonA2 may promote membrane partitioning and opted to analyze this hit further.

### PonA2 contributes to efficient mycobacterial growth following benzyl alcohol exposure

To test whether PonA2 contributes to benzyl alcohol tolerance and/or recovery, we constructed a clean deletion mutant (Δ*ponA2*) and treated with benzyl alcohol. We found that the number of colony-forming units (CFU) for Δ*ponA2* was comparable to that of wild-type (Fig. 3A and Supplementary Figure 1) immediately after benzyl alcohol treatment, but that Δ*ponA2* growth lagged during the early post-washout recovery period (Fig. 3B). The Δ*ponA2* growth delay was specific to benzyl alcohol as there was no delay after treatment with the vehicle control (DMSO). Furthermore, the defect in benzyl alcohol recovery was restored in a complemented strain (c*ponA2*), indicating that the defect is due to the lack of *ponA2*. These data suggest that PonA2 helps *M. smegmatis* to recover from membrane disruption.

**Figure 3.**
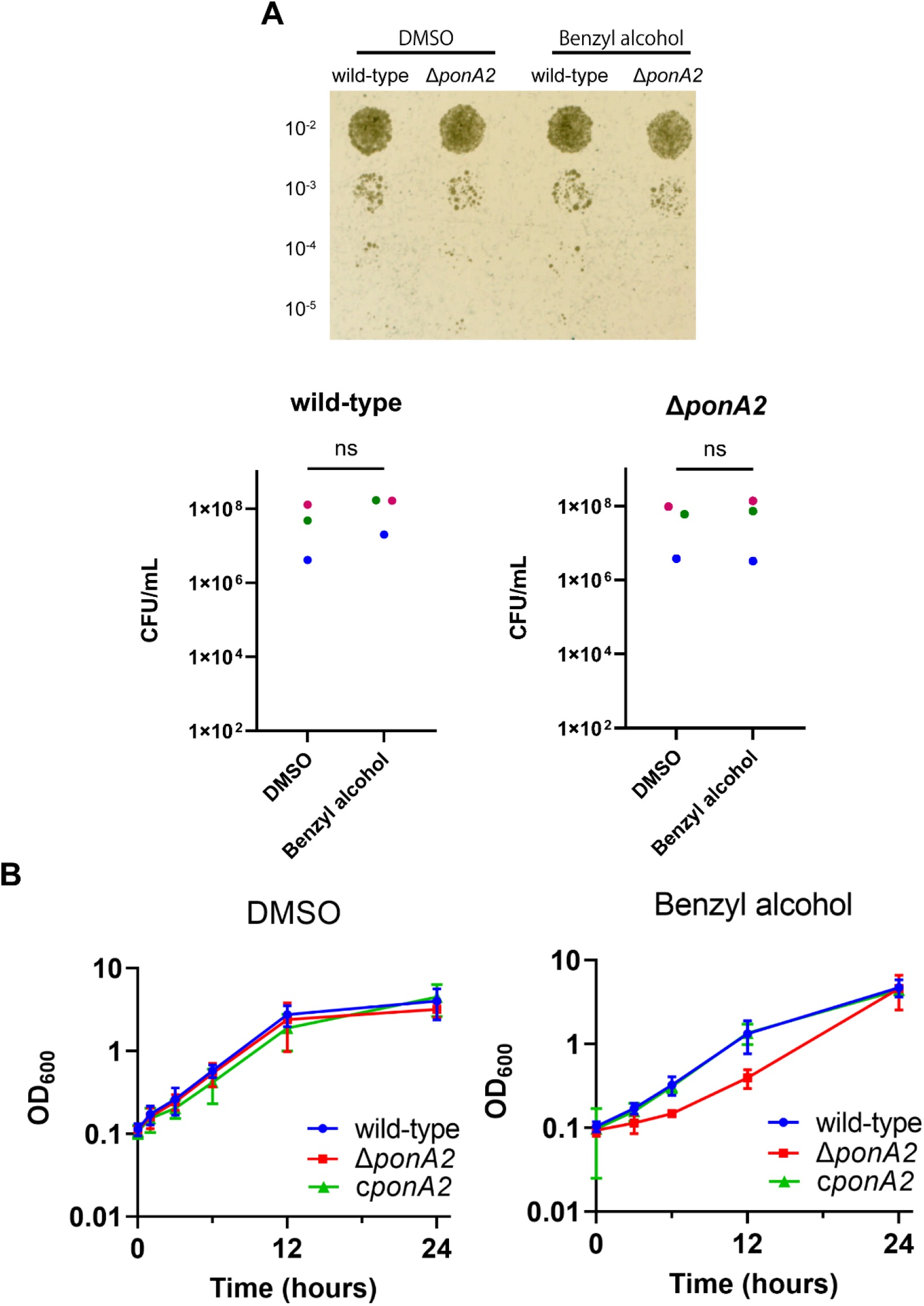
PonA2 contributes growth recovery from benzyl alcohol. (A) Top, wild-type or Δ*ponA2 M. smegmatis* were treated with benzyl alcohol for 1 hour, then 10-fold serial dilutions were spotted on Middlebrook 7H10 agar. Bottom, colony-forming units (CFU) were calculated from three biological replicates. Colors correspond to same-day replicates. ns, no statistically-significant difference. (B) Benzyl alcohol- or DMSO vehicle-treated wild-type, Δ*ponA2*, or complemented strain (c*ponA2*) were washed 3 times then grown in Middlebrook 7H9 medium. Data obtained from three independent experiments.

### PonA2 restores membrane partitioning after benzyl alcohol treatment

*M. smegmatis* eventually restores membrane partitioning after benzyl alcohol treatment (García-Heredia et al., 2021) (Fig. 1B) but the factors that promote repartitioning are unknown. We used subpolar enrichment of IMD-associated Ppm1 (Hayashi et al., 2016) as a readout for membrane partitioning before and after benzyl alcohol treatment. Prior to benzyl alcohol exposure, Ppm1 was enriched in the subpolar regions of wild-type, Δ*ponA2* and the complemented mutant (c*ponA2*; Fig. 4A and B). Immediately after benzyl alcohol exposure, subpolar enrichment was diminished for all three strains. During the recovery period, Ppm1 relocalized to the subpolar regions within ∼3 hours for wild-type and ∼1 hour for c*ponA2*. However, Ppm1 did not relocalize in Δ*ponA2* even up to 6 hours of outgrowth (Fig. 4A and C). These data indicate that PonA2 contributes to spatial repartitioning of the plasma membrane following benzyl alcohol-induced fluidization.

**Figure 4.**
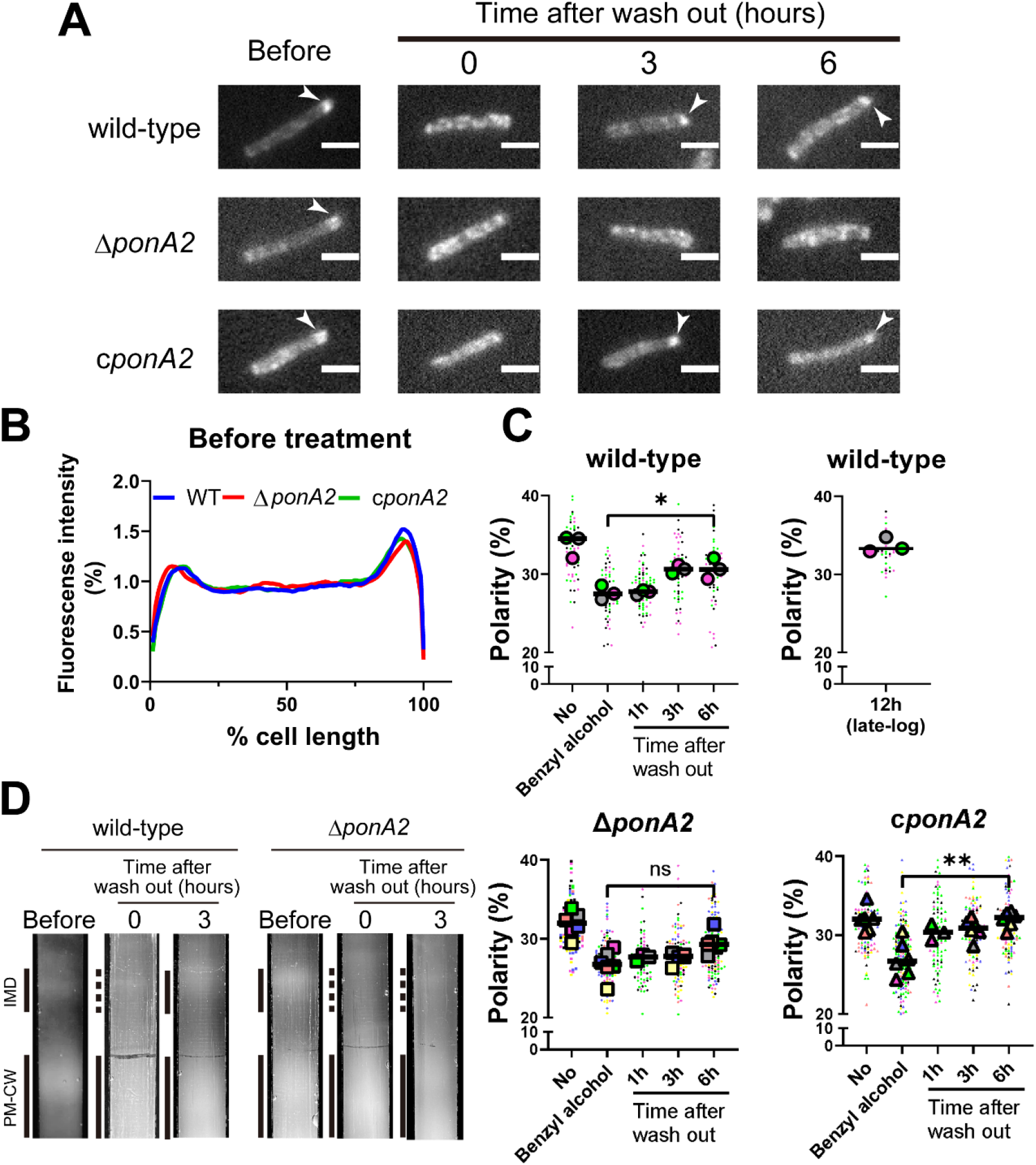
PonA2 promotes membrane repartitioning after benzyl alcohol treatment. (A) Fluorescence imaging of *M. smegmatis* expressing Ppm1-mNeonGreen before benzyl alcohol treatment or after benzyl alcohol washout. Arrowheads indicate subpolar foci of Ppm1-mNeonGreen. Scale bars, 2.5 µm. (B) Fluorescence of cells imaged as in (A) were quantitated from three independent experiments as in Fig. 1C. Lines show the average of total cells (50<n<69). (C) The percentage of signal associated with the distal 15% of rod-shaped cells are quantified to indicate polarity of fluorescence distribution. Each color represents an independent biological replicate. Smaller symbols are the means for bacteria in one imaging field and larger symbols are the means of the fields for a given independent replicate. Statistical significance was determined by the Kruskal–Wallis test, followed by Dunn’s multiple comparison test. ns, no statistically significant difference; *, p < 0.05; **, p < 0.01. (D) Lysates from *M. smegmatis* at indicated time points were sedimented in a sucrose density gradient.

As a complementary way to track membrane repartitioning, we examined the IMD and PM-CW biochemically. Under normal growth conditions, the IMD can be separated from the PM-CW by sucrose density gradient fractionation (García-Heredia et al., 2021; Hayashi et al., 2016, 2018; Morita et al., 2005); the IMD fractions are less dense and distinct from the PM-CW fractions. Benzyl alcohol treatment of both wild-type and Δ*ponA2 M. smegmatis* results in the apparent loss of membranous material from the IMD fractions (Fig. 4D) (García-Heredia et al., 2021). In contrast to wild-type, however, the mutant failed to recover the IMD within the 3 hours post-benzyl alcohol washout. This experiment suggests that, in addition to spatial repartitioning, PonA2 contributes to biochemical repartitioning of the plasma membrane post-benzyl alcohol.

We did not identify other enzymes involved in peptidoglycan biosynthesis from the Tn-seq analysis. Transposon insertions do not always interrupt gene function, and trans-complementation can occur when mutants are pooled. When tested individually, however, loss of other, non-essential cell wall synthases, including the monofunctional SEDS family transglycosylase RodA (Δ*rodA*) and multiple L,D-transpeptidases (Δ*ldtABE*) had no effect on *M. smegmatis* outgrowth post-benzyl alcohol (Supplementary Figure 2). These data suggest that promotion of membrane repartitioning is not a universal property of peptidoglycan biosynthetic enzymes.

### PonA2 does not affect membrane-cell wall interaction under basal conditions

Physical interactions between the plasma membrane and bilayer-extrinsic polymers have been proposed to maintain membrane partitioning in other systems (Daněk et al., 2020; Feraru et al., 2011; Martinière et al., 2012; McKenna et al., 2019; Wagner et al., 2020). We developed a microfluidics-based assay (Supplementary Fig. 3) to quantify and visualize membrane-cell wall interactions in *M. smegmatis*. In *Escherichia coli*, hyperosmotic shock causes severe plasmolysis, whereby the plasma membrane retracts from the cell wall, indicating that these structures are not strongly associated (Rojas et al., 2018). As plasmolysis occurs in areas with weak membrane-cell wall association, we predicted that plasmolysis bays in *M. smegmatis* would preferentially form at sites of IMD enrichment, *e.g.,* subpolar foci and midcell patches. We exposed wild-type and Δ*ponA2 M. smegmatis* in a microfluidics chamber to hyperosmotic shock and measured the number and location of plasmolysis bays. As expected, the location of bay formation correlated with the location of the IMD. However, there were no major differences in plasmolysis frequency or subcellular distribution between the two strains (Supplementary Fig. 3), suggesting that PonA2 likely does not promote membrane repartitioning via pre-existing, physical interactions between the plasma membrane and cell wall. This conjecture is further supported by our bulk biochemical assay (Fig. 4D), which demonstrates that the PM-CW (plasma membrane tightly associated with the cell wall) is isolable even in the absence of PonA2.

### PonA2 promotes membrane repartitioning and regrowth post-fluidization via its conserved transglycosylase domain

Given that PonA2 is a bifunctional transpeptidase/transglycosylase, we hypothesized that one or more of its enzymatic activities contributes to membrane homeostasis. Accordingly, we made point mutations to alter well-conserved, catalytically-active amino acids (Supplementary Figure 4), substitutions that have been previously shown to eliminate activity in *in vitro* assays for PBP1a (Born et al., 2006) and PBP1b (Terrak et al., 1999) in *E. coli*, and the transpeptidase function of PonA1 in *M. smegmatis* (Kieser et al., 2015b). Specifically, we complemented Δ*ponA2* with a *ponA2* allele that bears E108T (transglycosylase inactive (TG-)), S405A (transpeptidase inactive (TP-)), or both (TG-/TP-) mutations. Δ*ponA2* complemented with the TP-*ponA2* allele behaved similarly to Δ*ponA2* complemented with wild-type *ponA2* for both post-benzyl alcohol outgrowth (Fig. 5A) and membrane repartitioning (Fig. 5B, and 5C). However, the recovery of Δ*ponA2* complemented with the TG- or TP-/TG-*ponA2* allele was delayed in a manner comparable to uncomplemented Δ*ponA2*. These data suggest that PonA2 promotes membrane repartitioning and regrowth post-benzyl alcohol via its conserved transglycosylase domain.

**Figure 5.**
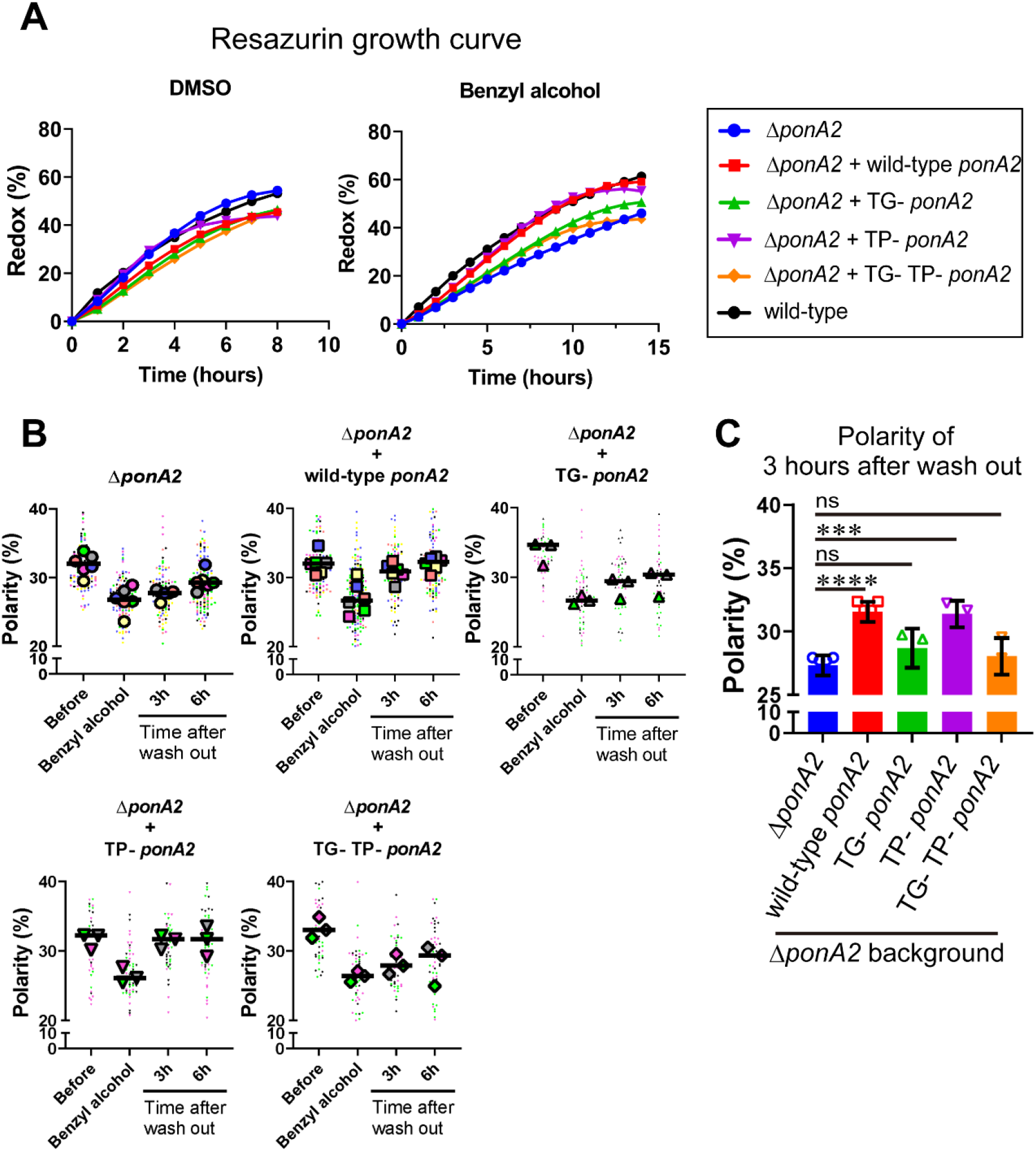
Conserved amino acid in the transglycosylase domain of PonA2 is required for efficient regrowth and membrane repartitioning post-benzyl alcohol. (A) Following exposure to benzyl alcohol or the DMSO vehicle control, cells were washed, resuspended in Middlebrook 7H9, and incubated with 0.0015% of resazurin. Data were obtained from three independent experiments and are means of biological duplicates or triplicates. (B) IMD marker polarities were assessed in mutants before, during, or 3- or 6-hr after benzyl alcohol treatment as in Fig. 4C. Data from Δ*ponA2* and Δ*ponA2* + wild-type *ponA2* are repeated from Figure 4C for ease of comparison. Each color represents an independent biological replicate. Smaller symbols are the means for bacteria in one imaging field and larger symbols are the means of the fields for a given independent replicate. (C) The polarity of 3-hour time points from Fig. 5B were compiled and compared across mutants. Statistical significance was determined by the Kruskal–Wallis test, followed by Dunn’s multiple comparison test. ns, no statistically significant difference; ****, p < 0.001; ***, p < 0.005.

### PonA2’s roles in supporting polar cell wall elongation, maintaining cell morphology and ensuring cell impermeability are genetically-separable from its role in membrane partitioning

We wondered whether PonA2 supports membrane homeostasis by localizing cell wall assembly. Previously we showed that the polarity of peptidoglycan synthesis decreases in the absence of RodA, a SEDS family transglycosylase (and upon treatment with moenomycin, an antibiotic that specifically inhibits transglycosylation of class A PBPs (aPBPs) such as PonA2 (Melzer et al., 2022). To more directly assay the function of PonA2, we labeled nascent peptidoglycan in wild-type and Δ*ponA2* after a brief incubation in the presence of alkyne-D-alanine-D-alanine (alkDADA, also called EDA-DA (García-Heredia et al., 2018; Liechti et al., 2014)) and detected the presence of the alkyne probe via copper-catalyzed alkyne-azide cycloaddition (CuAAC) to a fluorescent azide label. In wild-type and Δ*ponA2* mycobacteria, fluorescence was enriched at the cell poles, the sites of cell elongation in this genus (Aldridge et al., 2012; Joyce et al., 2012; Kieser and Rubin, 2014; Meniche et al., 2014; Santi et al., 2013; Singh et al., 2013; Thanky et al., 2007) and Fig. 6A). However, as with RodA absence or moenomycin treatment (Melzer et al., 2022) there was a modest but statistically significant decrease in the polarity of nascent peptidoglycan in Δ*ponA2* compared to wild-type (Fig. 6B and 6C). Moreover, a subpopulation of mutant cells had clear cell bulging (Fig. 6A and Supplementary Figure 5), a phenotype that has long been linked to peptidoglycan defects (Burdon, 1946; Chung et al., 2009; Hett et al., 2010; Huang et al., 2008; Typas et al., 2010; Vigouroux et al., 2020). The morphological defects are consistent with prior observations of aberrant width and morphology, respectively, in Δ*ponA2 M. tuberculosis* (Kieser et al., 2015a) and stationary phase Δ*ponA2 M. smegmatis* (Patru and Pavelka, 2010). However, in contrast to the role of PonA2 in post-benzyl alcohol membrane repartitioning and regrowth, which depend solely on its transglycosylation domain (Figs. 5A-5C), the role of PonA2 in localizing cell wall synthesis depended on both its transglycosylase and transpeptidase domains (Fig. 6D). Furthermore, the role of PonA2 in maintaining normal cell shape did not depend on either domain (Supplementary Figure 5). The genetic separability of the phenotypes suggests that PonA2’s contributions to membrane partitioning, polar peptidoglycan assembly and cell morphology are distinct.

**Figure 6.**
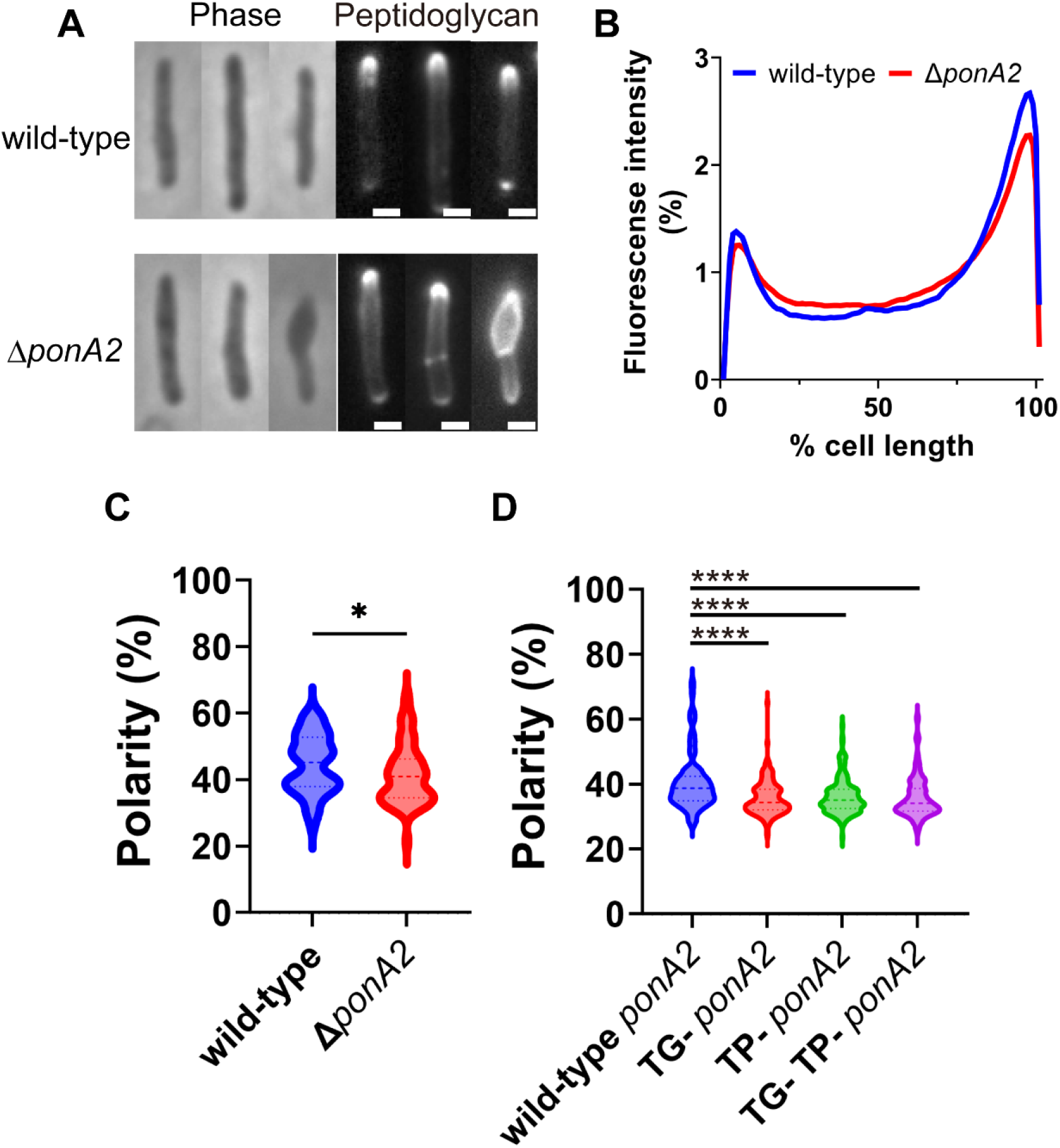
PonA2 localizes peptidoglycan synthesis and maintains cell morphology. (A) Nascent peptidoglycan in wild-type or Δ*ponA2 M. smegmatis* was labeled for 15 min (∼10% generation) with alkyne-D-alanine-D-alanine (alkDADA) followed by copper-catalyzed alkyne-azide cycloaddition (CuAAC) with AFDye488 Azide. Scale bars, 1 µm. (B) Wild-type and Δ*ponA2* strains were labeled as in (A) and subcellular fluorescence was quantitated as in Fig. 1C. Lines show the average of total cells (50 < n < 68). (C) The percentage of signal associated with the distal 15% of rod-shaped cells quantified to indicate polarity of fluorescence distribution. Mann-Whitney U p-Value, *, p < 0.05. (D) Polarity ratio of catalytically inactive mutants. Statistical significance was determined by the Kruskal–Wallis test, followed by Dunn’s multiple comparison test. ****, p < 0.001

Given that loss of PonA2 sensitizes mycobacteria to a variety of stresses (DeJesus et al., 2017a; Kieser et al., 2015a; Li et al., 2022; Patru and Pavelka, 2010; Vandal et al., 2008, 2009b), we also considered the possibility that the membrane partitioning and growth phenotypes of Δ*ponA2* uncovered by benzyl alcohol were associated with enhanced susceptibility to the chemical. For example, high fluidity can enhance the permeability of model membranes (Frallicciardi et al., 2022; Gabba et al., 2020; Lande et al., 1995; Rossignol et al., 1982). To test membrane permeability, we incubated wild-type and Δ*ponA2* with propidium iodide, a dye that fluoresces upon DNA intercalation and is normally membrane-impermeant. At baseline, neither wild-type nor Δ*ponA2* stained appreciably with propidium iodide (Supplementary Figure 6A). As expected, however, ∼20% of wild-type cells were propidium iodide-positive after benzyl alcohol treatment (Supplementary Figure 6A), indicating that benzyl alcohol-induced fluidization can compromise the membrane barrier. Moreover, ∼60% of the Δ*ponA2* cells were propidium iodide-positive (Supplementary Figure 6A). These data suggest that Δ*ponA2* cells are more permeable than wild-type following benzyl alcohol exposure.

Propidium iodide staining is often used to detect dead cells. However, the colony-forming units of bacteria with or without PonA2 or benzyl alcohol were similar (Fig. 3A), suggesting that enhanced permeability was not lethal. We reasoned that live, propidium iodide-positive cells may be able to grow upon dye washout, and that DNA synthesis and subsequent cell division would dilute the fluorescence over time. To test whether the propidium iodide-positive cells were alive, we examined the fluorescence of the propidium iodide-positive population after washing out the dye. As a negative control, we incubated heat-killed cells in growth medium and confirmed that there was no change in fluorescence over time. In contrast, the propidium iodide-positive populations of wild-type and Δ*ponA2* were reduced by half after three and six hours of outgrowth, respectively (Supplementary Figure 6B), a rate of dilution that roughly correlated with the bulk population growth rate of the strains during the same time frame (Fig. 3B. Thus, while loss of PonA2 exacerbates benzyl-alcohol-induced membrane permeabilization, the perturbations alone or combined do not compromise viability. Moreover, in contrast to the delayed growth and membrane repartitioning of benzyl alcohol-exposed Δ*ponA2*, enhanced membrane permeability was not complemented by the TP-*ponA2* allele (Supplementary Figure 6C). The genetic separation of these phenotypes implies that reinforcement of the membrane permeability barrier does not explain the role of PonA2 in membrane partitioning.

### De novo membrane partitioning is supported by the preexisting cell wall polymer but does not require concurrent cell wall polymerization

Efficient membrane repartitioning post-benzyl alcohol depends on the conserved transglycosylase domain of PonA2 (Fig. 5B and 5C). We previously demonstrated that early-stage inhibition of cell wall synthesis by D-cycloserine delocalizes IMD markers, but that it takes approximately one generation and the IMD remains biochemically isolable throughout (Hayashi 2018). We considered the possibility that active cell wall synthesis might initiate but not maintain membrane partitioning. However, when we treated wild-type *M. smegmatis* with different concentrations of D-cycloserine or aPBP transglycosylation inhibitor moenomycin during the recovery period after benzyl alcohol washout, we found that the growth-inhibitory effects of the antibiotics were similar regardless of whether the culture had been previously exposed to benzyl alcohol (Fig. 7A). These and our previous data imply that active cell wall polymerization is dispensable for membrane partitioning.

**Figure 7.**
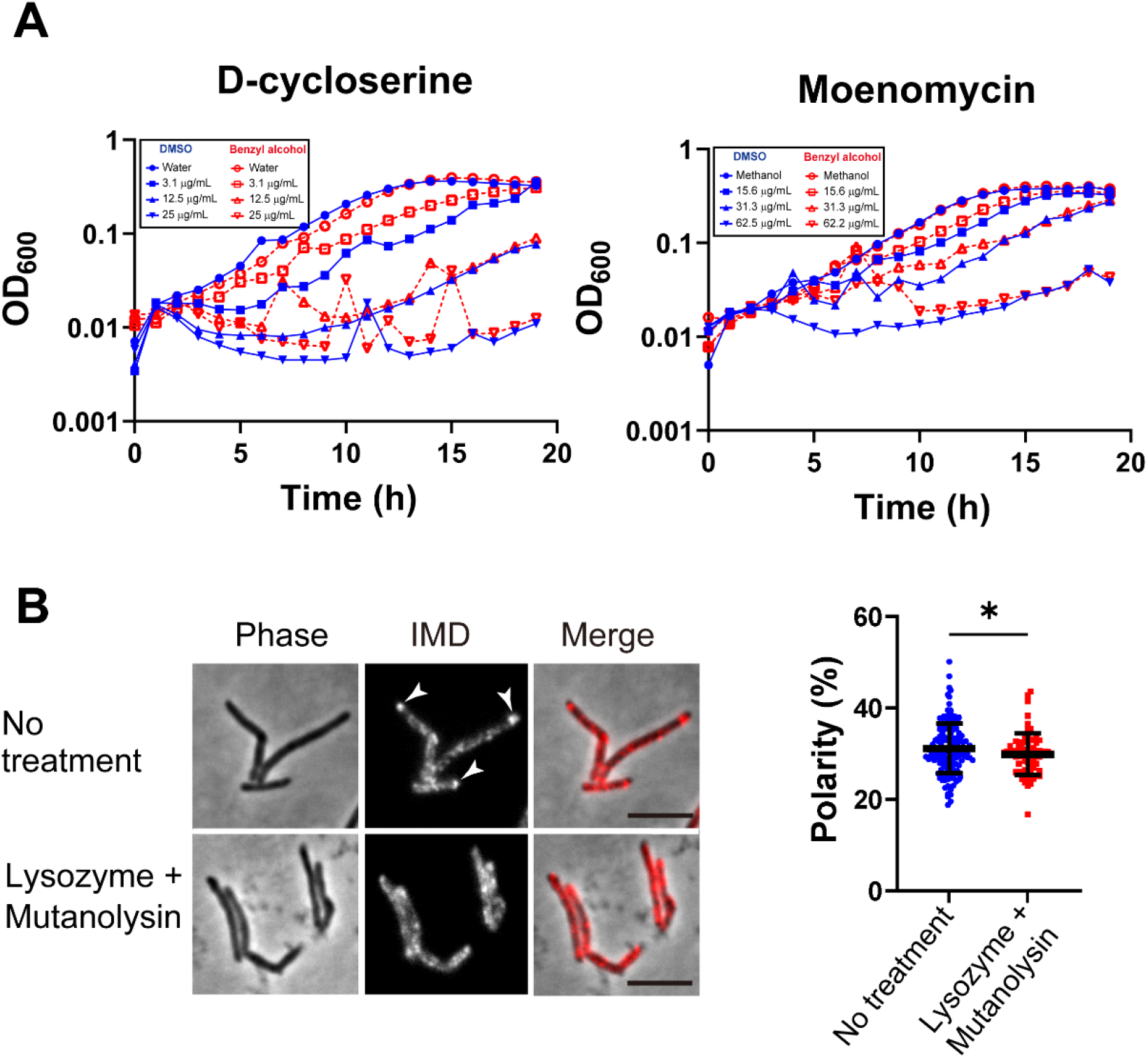
The integrity of the preexisting cell wall polymer, but not active cell wall polymerization, promotes membrane partitioning. (A) benzyl alcohol (red line)- or DMSO vehicle-(blue line) treated bacteria were washed then grown in Middlebrook 7H9 with D-cycloserine or moenomycin at the indicated concentrations. The lines show the average of 3 or 4 independent experiments. (B) Left, images of *M. smegmatis* expressing validated IMD marker mCherry-GlfT2 +/-60 min treatment with cell wall hydrolases lysozyme and mutanolysin. Arrowheads indicate subpolar foci of mCherry-GlfT2. Scale bars, 5 µm. Right, quantitation of mCherry-GlfT2 polarity for cells with no treatment (n = 67) or lysozyme/mutanolysin treatment (n = 151). The percentage of signal associated with the distal 15% of rod-shaped cells are quantified to indicate polarity of fluorescence distribution. Mann-Whitney U p-Value, *, p < 0.05.

We hypothesized that PonA2 transglycosylation promotes membrane repartitioning (Fig. 5B and 5C) by supporting cell wall integrity. While we previously showed that spheroplasting delocalizes multiple IMD markers (García-Heredia et al., 2021), complete removal of the cell wall is likely to have pleiotropic effects on membrane physiology. More recently, we showed that limited peptidoglycan digestion by the glycoside hydrolases lysozyme and mutanolysin delocalizes the IMD-enriched protein MurG (Melzer et al., 2022). To test the specificity of this observation, we examined the distribution of a second IMD marker, GlfT2 (Hayashi et al., 2016, 2018). As with MurG, we found that GlfT2 is rapidly delocalized upon lysozyme and mutanolysin treatment (Fig. 7B). Our results imply that membrane partitioning depends on the pre-existing cell wall polymer, rather than active cell wall polymerization.

## Discussion

Lateral organization is likely to be a key regulator of plasma membrane function yet is experimentally challenging to manipulate in living cells. Our inducible departitioning/repartioning model (Fig. 1) and subsequent screening (Fig. 2) is a generalizable approach to genetically dissect the mechanisms by which cellular membranes are partitioned.

Factors that establish and maintain plasma membrane partitioning are likely important for cell fitness but may be distinct from each other. We previously demonstrated a close correlation between membrane partitioning and cell growth in *M. smegmatis* (Hayashi et al., 2018) and that the known chemical fluidizer benzyl alcohol departitions the membrane and halts growth in this organism (García-Heredia et al., 2021). Building on these observations, we screened here for mycobacterial genes that counteract the growth-inhibitory effects of benzyl alcohol, a subset of which we hypothesized would also promote membrane partitioning. We identified a factor, cell wall synthase PonA2, that establishes membrane partitioning via its transglycosylase domain. Our earlier work had suggested that peptidoglycan damage and/or removal (García-Heredia et al., 2021; Melzer et al., 2022), but not inhibition of peptidoglycan synthesis (Hayashi et al., 2018), departition the *M. smegmatis* membrane. Similar, seemingly-incongruous observations have been made for the roles of the cell wall and its synthesis in other bacteria (Wagner et al., 2020) and in plants (Daněk et al., 2020). We resolve this conundrum for *M. smegmatis* by showing that active cell wall synthesis, including aPBP-mediated polymerization, is dispensable for efficient initiation of membrane partitioning, but that the pre-existing cell wall must be intact and properly polymerized with the help of PonA2.

In model membranes, the partitioning of bilayers supported on fabricated glycan networks can be controlled by glycan density (Subramaniam et al., 2013). While heterogenous networks stabilize membrane domains, homogenous networks suppresse macroscopic phase separation. In laterally growing, rod-shaped bacteria, it is thought that the SEDS-family transglycosylase RodA lays the template for cell wall elongation while the bifunctional, transglycosylase/transpeptidase aPBPs fill in the gaps for maintenance and repair (Cho et al., 2016; Mueller et al., 2019; Murphy et al., 2021; Paradis-Bleau et al., 2010; Typas et al., 2010; Vigouroux et al., 2020). Unlike the organisms in which this model has been tested, pole-growing *Mycobacteriales* lack the cytoskeletal protein MreB and do not require RodA for viability or shape (Arora et al., 2018) so the division of labor is less clear (Melzer et al., 2022; Sher et al., 2021). Given that PonA2 but not RodA contributes to membrane partitioning in *M. smegmatis*, PonA2 may regulate the 3-dimensional density and/or heterogeneity of the polymeric cell wall network.

We envision at least two scenarios by which the PonA2-built cell wall polymer regulates membrane partitioning. Membrane-bound proteins and/or other biomolecules may partition the bilayer by tethering it to the cell wall and influencing protein and lipid diffusion. In this first model, PonA2 promotes membrane partitioning by creating a cell wall structure conducive to tethering. While PonA2 does not affect membrane-cell wall interactions in bulk or under basal conditions (Fig. 4D and Supplementary Fig. 3), we cannot currently rule out subtle changes in re-tethering during post-benzyl alcohol recovery. Alternatively, in the second model, the cell wall polymer interacts with the membrane bilayer in a more direct fashion. For example, the partitioning behavior of model membranes in response to fabricated glycan networks (and in the absence of potential tethering molecules) (Subramaniam et al., 2013) was proposed to be the result of changes in liquid-ordered phase stability or in the line tension between liquid-ordered and liquid-disordered phases.

PonA2 is not required for *M. smegmatis* or *M. tuberculosis* growth. However, the bifunctional transglycosylase/transpeptidase protects these organisms from various stresses, including heat, a classic membrane fluidizer (Patru and Pavelka, 2010). Loss of *ponA2* also sensitizes mycobacteria to antibiotics with different structures and cellular targets (Kieser et al., 2015a; Li et al., 2022; Patru and Pavelka, 2010; Vandal et al., 2008, 2009a), a phenotype that suggests enhanced small molecule permeability. Aberrant peptidoglycan synthesis in the absence of PonA2 may disrupt the cell envelope layers outside of the peptidoglycan, including the mycomembrane. As mycomembrane-disrupting mutations can sensitize mycobacteria to many antibiotics (Gao et al., 2003; Li et al., 2022; Liu and Nikaido, 1999; Nguyen et al., 2005; Philalay et al., 2004; Vilchèze et al., 2014), it is often assumed that the mycomembrane is the primary determinant of mycobacterial impermeability and intrinsic antibiotic resistance. Our work suggests that *ponA2* mutations can also impact the organization and integrity of the layer inside of the peptidoglycan, the plasma membrane. It is an open question how plasma membrane defects contribute to the stress-specific phenotypes of mycobacterial *ponA2* mutants.

We found that PonA2’s conserved transpeptidase and transglycosylase activities are both necessary for the localization of cell wall synthesis and maintenance of impermeability. In contrast, neither activity is needed for the maintenance of cell morphology, and only the conserved transglycosylase activity contributes to membrane partitioning and growth after membrane fluidization. The roles of these individual activities in PonA2-mediated stress protection have not been reported. However, differential requirements for individual enzymatic functions have been characterized for the closely-related aPBP PonA1. PonA1’s conserved transglycosylase domain is critical for viability and normal cell length in *M. smegmatis* and normal production of phthiocerol dimycocerosate (PDIM) in *M. tuberculosis*; and both domains are needed for maintaining correct cell length in *M. tuberculosis* (Kieser et al., 2015b). The potential association between PonA1 and the virulence-associated lipid PDIM (Camacho et al., 1999, 2001; Cox et al., 1999; Rens et al., 2021) is particularly intriguing given that the biosynthesis of PDIM is partitioned within the IMD of the *M. tuberculosis* plasma membrane (Puffal et al., 2022). We previously found that perturbations that departition the membrane can negatively impact the efficiency and precision of pathways embedded therein (García-Heredia et al., 2021). Given that loss of PonA2 sensitizes *M. tuberculosis* to host antimicrobial responses (Smith et al., 2022; Vandal et al., 2009a; Wagner et al., 2020), it is possible that promotion of membrane partitioning by PonA2 transglycosylation (Fig. 5B and 5C) indirectly supports robust PDIM biosynthesis.

Assembly of the cell wall occurs adjacent to, and using precursors from, the plasma membrane. We show here that the completed cell wall polymer partitions the *M. smegmatis* plasma membrane into regions that are spatially (Fig. 5B), biochemically (Fig. 5D), and biophysically (Supplementary Fig. 3) distinct. At the same time, individual steps of the peptidoglycan synthesis pathway are partitioned within bacterial membrane. For example, regions of increased fluidity (RIFs) are enriched for MurG, the synthase for the polyprenol phosphate-linked cell wall precursor lipid II (Müller et al., 2016; Strahl et al., 2014). Functional membrane microdomains (FMMs) are enriched for lipid II flippase MurJ and for extracellular synthases that use lipid II to assemble the cell wall (García-Fernández et al., 2017). In mycobacteria, we have demonstrated that lipid II is made in the IMD, then trafficked to, and likely polymerized in, the PM-CW (García-Heredia et al., 2021). Our data now suggest that the positive feedback loop between membrane and cell wall organization initiates with the cell wall polymer. Damage to the cell wall, which we accomplish experimentally by treatment with glycoside hydrolases, interrupts this pro-growth feedback loop and departitions the membrane (Fig. 7B and Fig. 8, (García-Heredia et al., 2021; Melzer et al., 2022), which in turn delocalizes the synthesis of peptidoglycan and other cell envelope components to the sidewall (García-Heredia et al., 2018, 2021). We hypothesize that delocalized envelope synthesis enables mycobacteria to robustly respond to and repair cell-wide damage (García-Heredia et al., 2018). Once the stress has passed, we further posit that the pro-growth, membrane-cell wall feedback loop begins anew. The interplay between membrane and cell wall organization may enable mycobacterial cells to adjustpolar growth and sidewall repair as needed.

**Figure 8.**
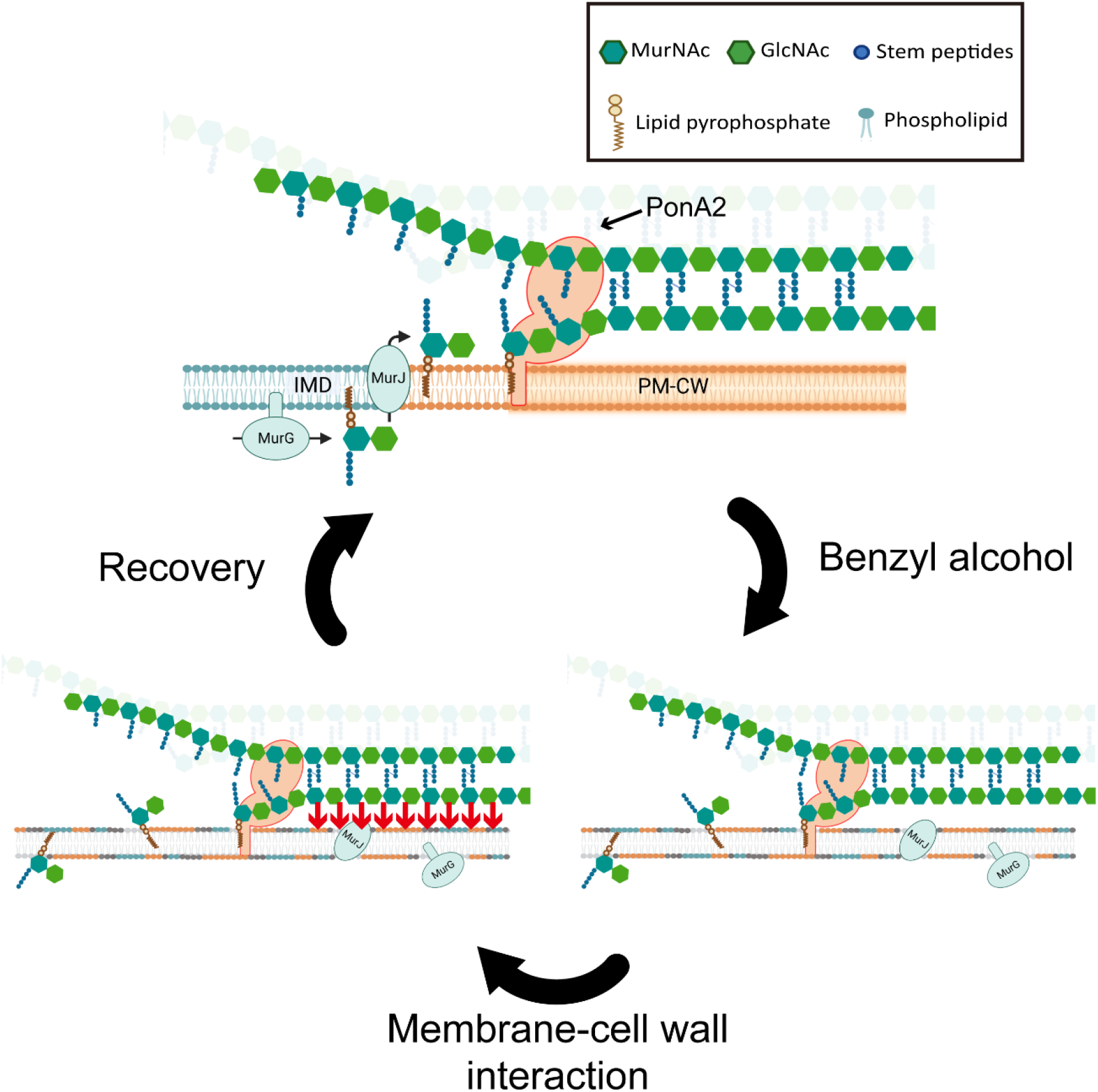
Working model for PonA2 in mycobacterial plasma membrane organization. PonA2, likely enriched in the PM-CW, polymerizes peptidoglycan using lipid II as a donor substrate, which in turn is produced by MurG in the IMD. PonA2 also cross-links nascent glycan strands into the existing cell wall. Upon membrane fluidization, lipid II and IMD-enriched proteins are no longer confined in the IMD (Garcia-Heredia 2021). Reestablishment of the IMD and growth recovery after membrane fluidization depend on the pre-existing, PonA2-built cell wall polymer. We speculate that PonA2-built peptidoglycan directly or indirectly interacts with the plasma membrane (red arrows; see Discussion for details) to promote repartitioning of the membrane into the IMD and PM-CW.

## Acknowledgement

We thank Drs. Eric Rubin and Chidi Akusobi for Tn-seq guidance, Alam García-Heredia for advice, and Jungwoo Lee and Jun-Goo Kwak for plate reader guidance. Some of the data were obtained at the University of Massachusetts Flow Cytometry, Biophysical Characterization, and Genomics Resource Core Facilities, with support from the Institute for Applied Life Sciences and directors Drs. Amy Burnside, Stephen Eyles and Ravi Ranjan. Research was supported by funds from the National Institutes of Health (NIH) under awards R21 AI144748 (YSM and MSS), DP2 AI138238 (MSS), R03 AI140259-01 (YSM), R35GM143057 (ERR), and T32 GM008515 (ESM, under the Chemistry and Biology Interface Program at the University of Massachusetts Amherst). TK was supported by a postdoctoral fellowship from Uehara Memorial Foundation.

## Declaration of interests

The authors declare no competing interests.

## Materials and Methods

### Bacterial strains and growth conditions

Markerless, knock-in *M. smegmatis* strains expressing both HA-mCherry-GlfT2 and Ppm1-mNeonGreen-cMyc or HA-mCherry-GlfT2 alone were previously established (Hayashi et al., 2016). *Mycobacterium smegmatis* mc^2^155 (wild-type), Δ*ponA2*, and Δ*ponA2 L5*::*ponA2* (wild-type and various alleles of *ponA2*) were grown in Middlebrook 7H9 growth medium (BD Difco, Franklin Lakes, NJ) supplemented with 0.4% (vol/vol) glycerol, and 0.05% (vol/vol) Tween-80 (Sigma–Aldrich, St. Louis, MO), as well as kanamycin (50 μg/mL) and/or hygromycin (50 μg/mL) where appropriate. Bacteria were grown at 37 °C with shaking at 130 rpm. For chemical treatments, stocks of 5 M benzyl alcohol in DMSO (Sigma–Aldrich), 0.2 M dibucaine (Sigma– Aldrich), 10% SDS (MP Biomedical), or undiluted oleic acid (Acros Organics) were respectively added to final concentrations of 100 mM, 0.5 mM, 0.05%, and 1.5 mg/mL to log-phase cultures. Phosphate-buffered saline (PBS) with 0.05% (vol/vol) Tween-80 (PBST) was used to wash out benzyl alcohol prior to resuspending bacteria in Middlebrook 7H9.

### Construction of plasmids and mutants

pMUM264 — To delete the endogenous *ponA2* gene, we amplified upstream and downstream regions of *ponA2* using the primers shown in Supplementary Table. These 2 fragments were assembled into pCOM1 (Hayashi et al., 2016) at Van91I sites by Gibson assembly (New England Biolabs). The assembled plasmid, pMUM264, was transformed into *M. smegmatis* by electroporation, and positive clones were isolated based on hygromycin resistance and SacB-dependent sucrose sensitivity. Correct deletion of the *ponA2* gene was confirmed by PCR.

pMUM280 — Primers A980 and A981 were designed to amplify *ponA2*, including 192 bp of upstream native promoter region from wild-type. The PCR fragment was assembled into pMUM 126 (Hayashi et al., 2016) at KpnI-XbaI sites by Gibson assembly®. The assembled plasmid, pMUM280, was transformed into *M. smegmatis* by electroporation, and positive clones were isolated based on kanamycin resistance.

pMUM293 (TG-) and 294 (TP-) — Primers A995 to A998 were designed to make point mutations of *ponA2* as shown in Supplementary Table using the Q5 Site-Directed Mutagenesis Kit (New England Biolabs). After mutations were confirmed by sequencing, the resulting plasmid, pMUM293 or 294, was transformed into *M. smegmatis* as above.

pMUM295 (TG-/TP-) — The part of pMUM293 which includes the TG region was digested by SacI and MluI and the fragment was inserted into the same region of pMUM294 by ligation.

### Transposon library construction

An transposon library of *M. smegmatis* were made by using Himar mutagenesis as previously described (Long et al., 2015; Siegrist and Rubin, 2009). Briefly, ΦMycoMarT7 phage (Piddock et al., 2000) and a log-phase *M. smegmatis* culture were mixed and incubated for at 37 °C for 4 h. Cells are spread on Middlebrook 7H10 medium supplemented with 0.5% glycerol, 0.05% Tween 80, and 50 μg/mL of kanamycin and incubated for 2-3 days at 37 °C, yielding a library of ∼10^5^ mutants. The library was harvested by scraping and stored as frozen stocks in Middlebrook 7H9 medium with 25% glycerol at 80 °C for further experiments. Library coverage of TA dinucleotide sites was determined to be ∼35% by Illumina sequencing.

### Benzyl alcohol selection of transposon libraries

A frozen stock was thawed and 20 μL of the stock was inoculated to 20 mL of Middlebrook 7H9 medium. After overnight incubation at 37 °C to allow the library to recover, this library starter culture was subcultured into 100 mL cultures to make a log-phase culture. The log-phase culture was treated by benzyl alcohol or DMSO vehicle control for one hour. Both cultures were washed 3 times with PBST and resuspended in Middlebrook 7H9 at a starting OD_600_ of 0.01. The cultures were incubated at 37 °C until the OD_600_ was 1.0 in order to standardize the number of outgrowth generations between libraries to approximately 6.5 generations.

### Sequencing of transposon mutant libraries

Genomic DNA was extracted from benzyl alcohol- or DMSO-treated transposon libraries, and the library mutant composition was determined by sequencing amplicons of the transposon-genome junctions as previously described using primers indicated in Supplementary Table (DeJesus et al., 2017b; Long et al., 2015). On average, library sequencing yielded between 0.5 million and 4 million unique transposon-inserted-sequences which cover over 35% of the possible TA sites in the genome.

### Mapping and quantification of transposon insertions

Raw sequence data were processed using the TPP tool from the TRANSIT TnSeq analysis platform (DeJesus et al., 2015), and transposon genome junctions were mapped to the *M. smegmatis* mc^2^155 reference genome (GenBank accession number NC_018143.1) using the Burroughs-Wheeler aligner (Li and Durbin, 2009). To account for possible PCR amplification biases, reads corresponding to the same TA site and possessing the same 7-nucleotide barcode were derived from the same template, and these duplicate reads were discarded from the final template counts. Data in Fig.2 were obtained from 3 independent experiments.

### Identification of genes affecting fitness under antibiotic selection

Genes conditionally affecting fitness in the presence of benzyl alcohol were identified using the resampling test module in the TRANSIT analysis platform as previously described (DeJesus et al., 2015, 2017b). In brief, we treated DMSO (a control treatment) and benzyl alcohol to transposon mutant library in triplicate. After washing out by PBST thrice, OD_600_ was adjusted to 0.01. The cultures were incubated for 16-24 hours to OD_600_ ∼1.0. DNA was isolated from 30 mL of culture, sequenced, and analyzed as described in the previous work (DeJesus et al., 2015, 2017b).

### CFUs and growth curves

Wildtype, Δ*ponA2*, and c*ponA2* cells were grown to stationary phase, then back-diluted and allowed to grow overnight to log phase (OD_600_ 0.5-0.8). Cultures were treated with DMSO or benzyl alcohol (100 mM of final concentration). Cultures were incubated at 37°C shaking at 150 rpm for 1 hour. The treated cultures were washed with PBST for 3 times and resuspended in Middlebrook 7H9 medium at a starting OD_600_ of 0.1 for continuous OD measurement in 125 mL flasks (Fig. 3C) or 96-well-plate with antibiotics (Fig. 7A). Biotek Synergy 2 was used for the growth curve in 96-well-plate. Aliquots (20 μL) were serial diluted with 7H9 media (200 μL) and 5 μL of aliquot is plated for colony-forming units (CFU).

### Resazurin growth curves

This was performed as described in the previous literature (Eagen et al., 2018). In brief, cells were washed three times with PBST and resuspended in Middlebrook 7H9 following a 1-hour treatment with DMSO vehicle control or benzyl alcohol. 200 µL of culture and 20 µL of 0.015% (w/v) resazurin (Acros Organics) were mixed in 96-well-plates and absorbance at 570 nm and 600 nm were measured by the Synergy H1 Hybrid microplate reader (BioTek) overnight. Percent of reduced resazurin was calculated as before (Eagen et al., 2018).

### Microscopy and image analysis

Bacterial culture was dropped onto the glass plate with agar pad (1% agarose in water). Images were acquired on Nikon Eclipse E600. Cell outlines were traced using Oufti (Nguyen et al., 2007; Paintdakhi et al., 2016). Intensity profiles were generated using MATLAB code as described in (Vandal et al., 2009b). Polarity ratios were calculated by combining signal values for 15% of the cell length on either pole and dividing the sum by total cell fluorescence. Super plots were generated as described (Lord et al., 2020).

### Membrane fractionation

Log-phase *M. smegmatis* treated or not with benzyl alcohol were harvested by centrifugation and washed in PBST. One gram of wet pellet was resuspended in 4 mL of lysis buffer containing 25 mM HEPES (pH 7.4), 20% (wt/vol) sucrose, 2 mM EGTA, and a protease inhibitor cocktail (ThermoFisher Scientific, Waltham, MA) as described (Morita et al., 2005). Bacteria were lysed and fractionated as in the previous literature (García-Heredia et al., 2021, 2021; Hayashi et al., 2016; Morita et al., 2005).

### Cell wall damage

Cells expressing mCherry-GlfT2 were grown to stationary phase, then back diluted and allowed to grow overnight to log phase (OD_600_ = 0.5-0.8). Lysozyme (Sigma-Aldrich) was dissolved and filter-sterilized freshly. The treatment condition was at final concentration of 500 μg/mL of lysozyme and 500 U/mL of Mutanolysin (Sigma-Aldrich). Cultures were incubated at 37°C shaking at 300 rpm in Benchmark Scientific MultiTherm Shaker H5000-H for 1 hour. Cells were imaged as described above.

### Cell envelope labeling

AlkDADA was custom synthesized by WuXi Apptec. Mid-log *M. smegmatis* was labeled with 2 mM alkDADA for 15 min. Cells were washed with PBST containing 0.01% BSA (PBSTB), and fixed in 2% formaldehyde at room temperature for 10 minutes. Cells were washed twice and applied for the reaction with CuAAC AFDye488 Azide (Click Chemistry Tool, Scottsdale, AZ) as described (García-Heredia et al., 2018; Siegrist et al., 2013).

### Cell width morphology profiles

Cells were placed on an agar pad slide, and imaged by phase microscopy (Nikon Eclipse E600, Nikon Eclipse Ti with 100x objectives). From the phase microscopy images, cells were outlined and segmented using Oufti (Paintdakhi et al., 2016). Cell width data were exported from Oufti and analyzed using a custom Python script. Using this script, cell lengths were normalized to a length of 1 (midcell = 0.5) and their width along their length was plotted as a line with each line representing a single cell. Multiple cell width profiles were superimposed on top of each other to visualize the major morphological trend (rod vs. blebbed). Additionally, percentages of cells with maximum widths greater than or equal to 0.95 µm (green dotted line) were counted and the total percentage of cells obtaining widths at or above these thresholds were displayed.

### Imaging in microfluidic devices

We used a Nikon Eclipse Ti2-E inverted fluorescence microscope with a 100× (NA 1.40) oil-immersion objective for imaging DU885 electron-multiplying charge-coupled device camera (Andor) for imaging (Rojas et al., 2018). Devices sere kept at 37 °C for imaging. Cells were streaked out on LB agar containing 50 µg/mL hygromycin and incubated at 37°C for 2-3 days. Single colonies were inoculated into Middlebrook 7H9 containing 50 µg/mL hygromycin and incubated for ∼48 hours at 37°C. The cells were then back diluted and added to the B04A microfluidic perfusion plate (CellASIC) during exponential phase. Plates were loaded with medium pre-warmed to 37°C. Cells were loaded into the plate, which was incubated at 37°C, without shaking, for 30 min before imaging. Medium was exchanged using the ONIX microfluidic platform (CellASIC). In the case where cells were stained with RADA, a TAMRA-based fluorescent D-amino acid (TOCRIS), 1 µM RADA was added to the culture upon back dilution. If the cells were not stained with RADA, Alexa Fluor 647 NHS succinimidyl ester (Thermo Fisher Scientific) was added to the media as an occlusion dye (it is not cell wall permeable and thus can be used to track the cells). Cells were perfused with Middlebrook 7H9 medium for 5 minutes and then hyperosmotically shocked with 7H9 + 3 M sorbitol for 10 minutes.

## Supplementary Information

**Supplementary Figure 1.**
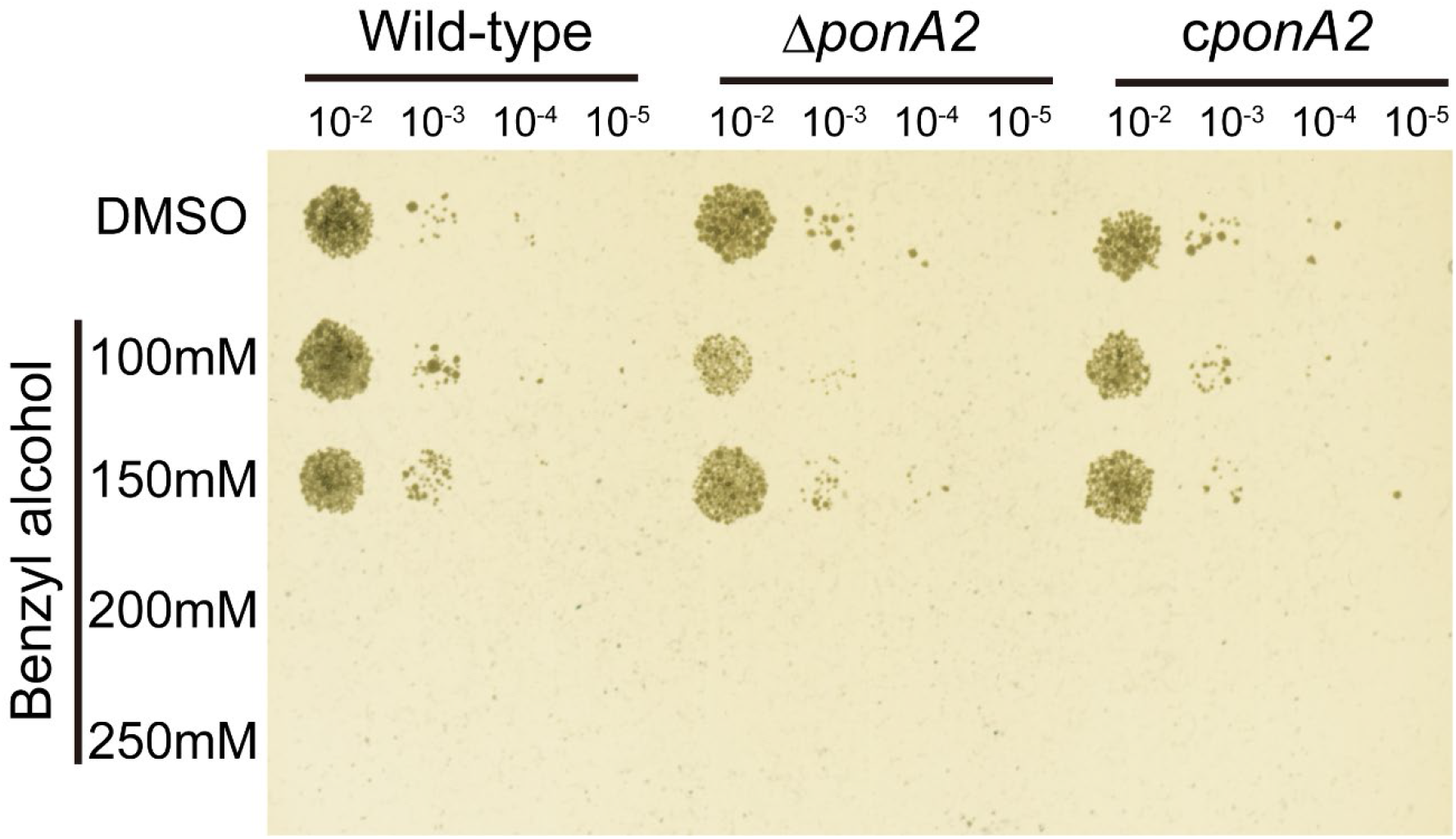
PonA2 is dispensable for survival during benzyl alcohol treatment. Wild type, Δ*ponA2*, or complemented *M. smegmatis* (c*ponA2*) were treated with benzyl alcohol at indicated concentration for 1 hour, and then 10-fold serial dilutions were spotted on Middlebrook 7H10 agar.

**Supplementary Figure 2.**
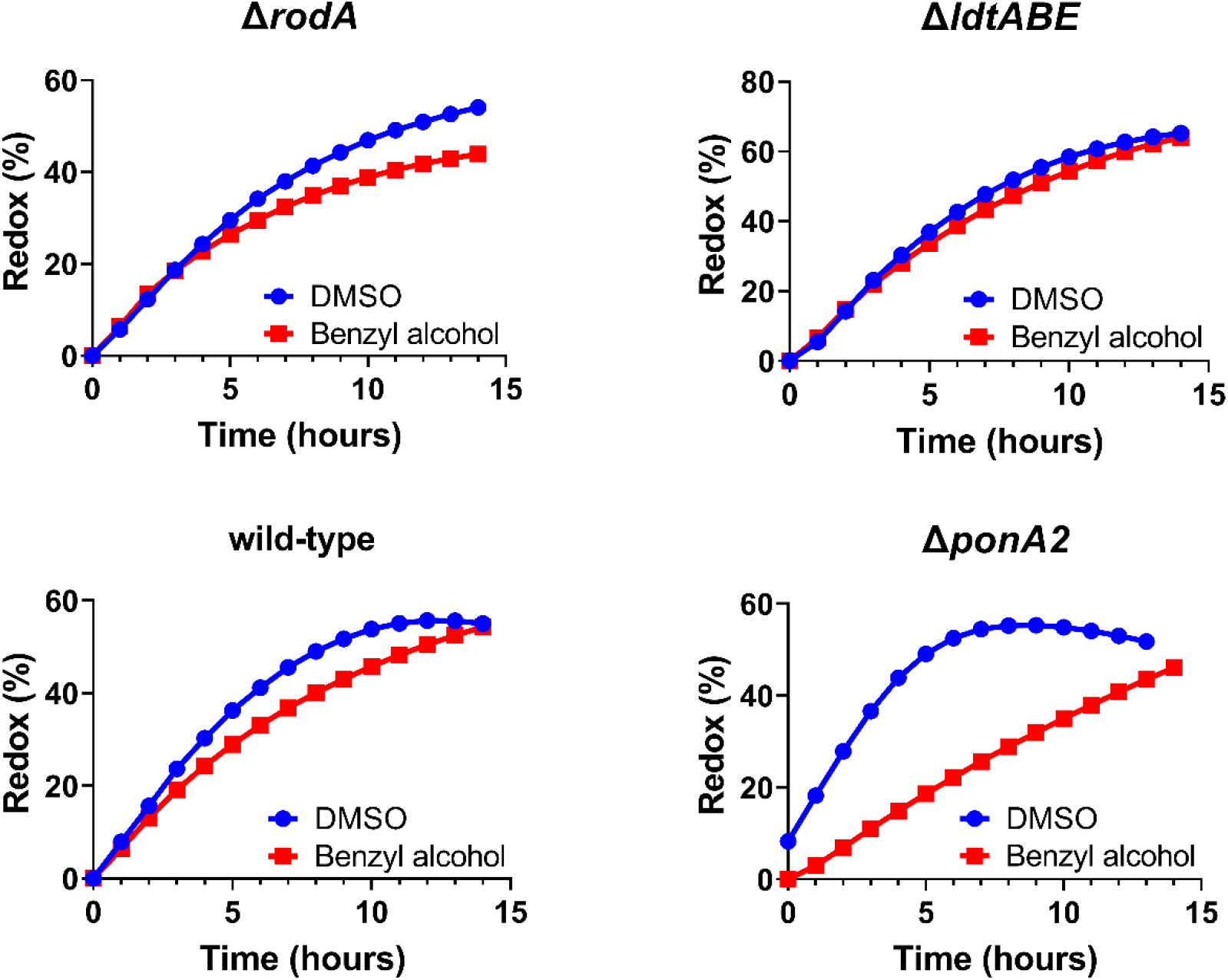
Following exposure to benzyl alcohol or the DMSO vehicle control, cells were washed, resuspended in Middlebrook 7H9, and incubated with 0.0015% of resazurin. Data were obtained from three independent experiments and are means of biological duplicates or triplicates. Data for wild-type and Δ*ponA2* are reproduced from Fig. 5 for comparison.

**Supplementary Figure 3.**
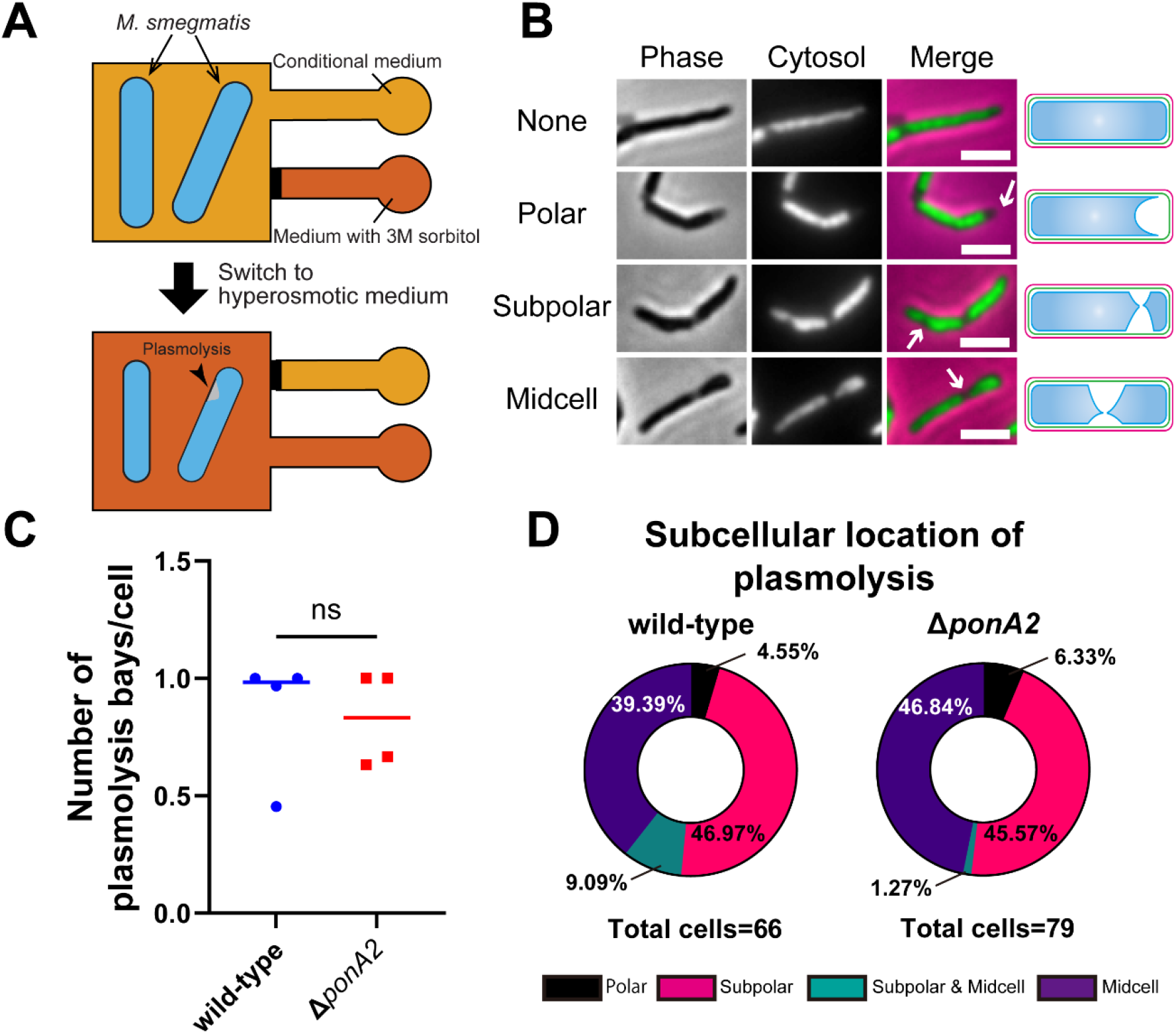
Bacteria were subjected to a large hyperosmotic shock, which depletes hydrostatic turgor pressure. High-resolution microscopy was then used to monitor the effects of turgor depletion on subcellular morphology. (A) Schematic of the microfluidics plasmolysis assay. GFP-expressing *M. smegmatis* are subjected to hyperosmotic shock (3 M sorbitol) in microfluidics device and imaged before, during and after shock. (B) Cytoplasmic GFP enables visualization and quantitation of plasmolysis bays. Arrows indicate representative sites of plasmolysis. Scale bars, 2.5 μm. (C) Plasmolysis bays/cell were calculated for wild-type and Δ*ponA2 M. smegmatis* from four microcolonies from two independent experiments. ns, not statistically significant by Mann-Whitney U test. (D) Analysis of the sites of plasmolysis (as defined in panel B) from the same experiment as (C). n = 113 for wild-type and 93 for Δ*ponA2 M. smegmatis*. The distribution of plasmolysis bays in the two strains was not statistically significant by Chi-square test.

**Supplementary Figure 4.**
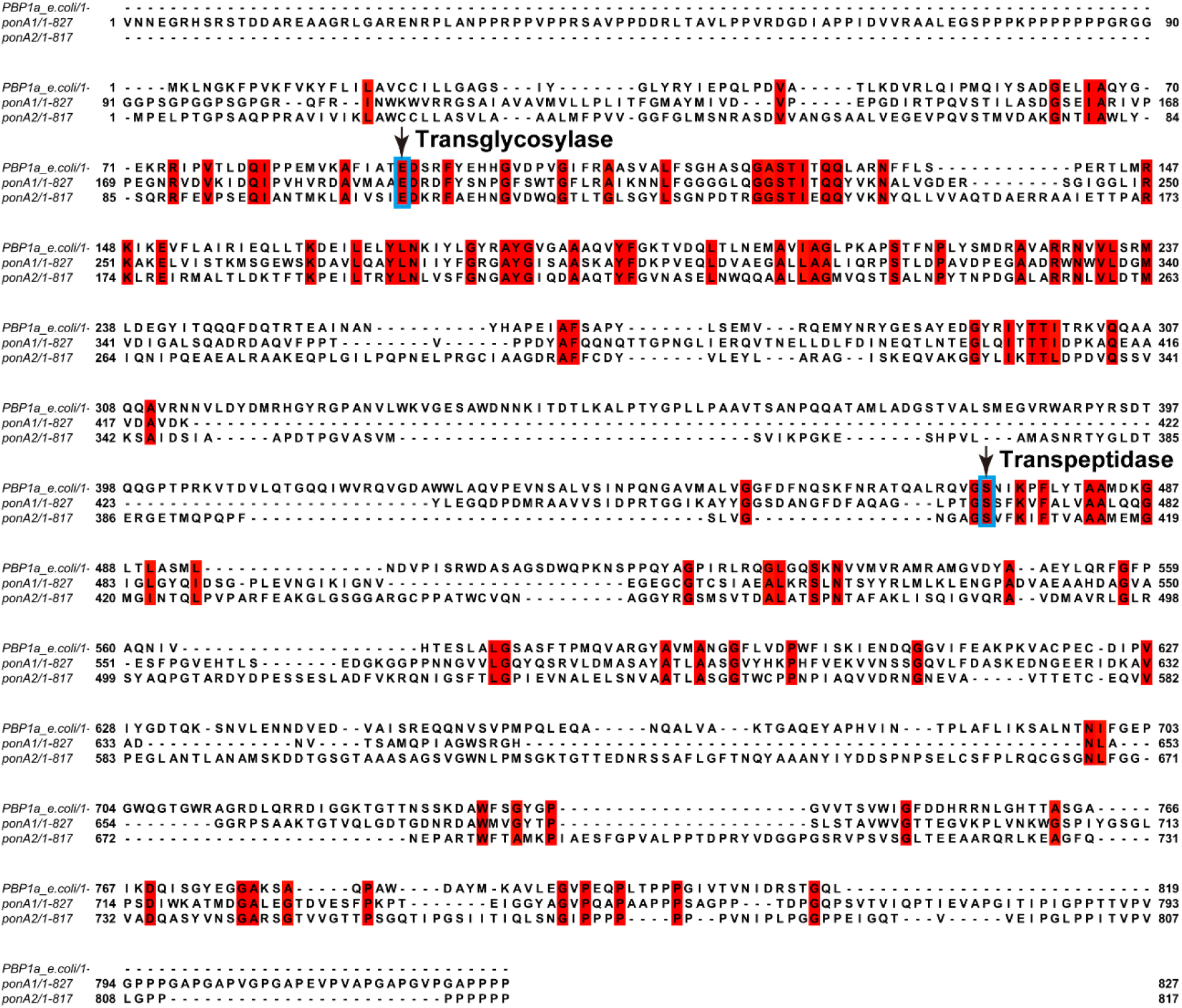
Protein alignments of PBP1a of *E. coli* and PonA1 and PonA2 of *M. smegmatis*. As in *M. smegmatis ponA1* (Kieser, et al., 2015), the true start codon of *ponA2* is located in the upstream of the annotation in MycoBrowser (Kapopoulou et al., 2011). That is, *M. smegmatis ponA2* starts from 87 base pair upstream of MSMEG_6201. Red shades indicate conserved amino acids. Arrows and blue squares indicate the amino acids that were mutated. Amino acid sequences were aligned using Jalview2 (Waterhouse et al. 2009).

**Supplementary Figure 5.**
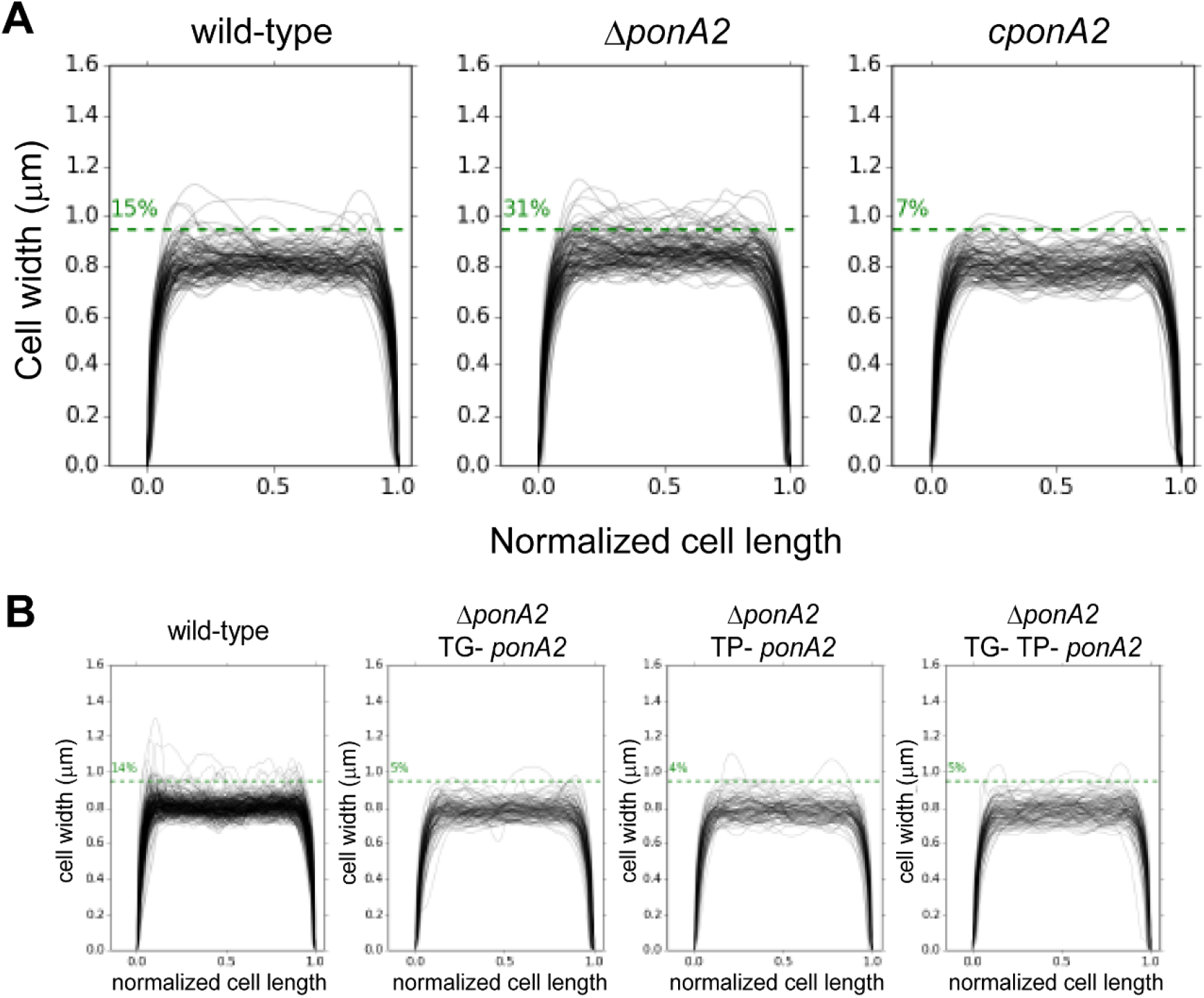
Cell width profiles of cells along the normalized cell length were determined by Oufti. (A) Cell widths of wild-type (n = 116), Δ*ponA2* (n = 124), and c*ponA2* (n = 107). (B) Cell widths of wild-type (n = 278), Δ*ponA2* TG-*ponA2* (n = 120), Δ*ponA2* TP-*ponA2* (n = 101) and Δ*ponA2* TG- TP- *ponA2* (n = 109) were quantified from three independent experiment. Cell lengths were normalized.

**Supplementary Figure 6.**
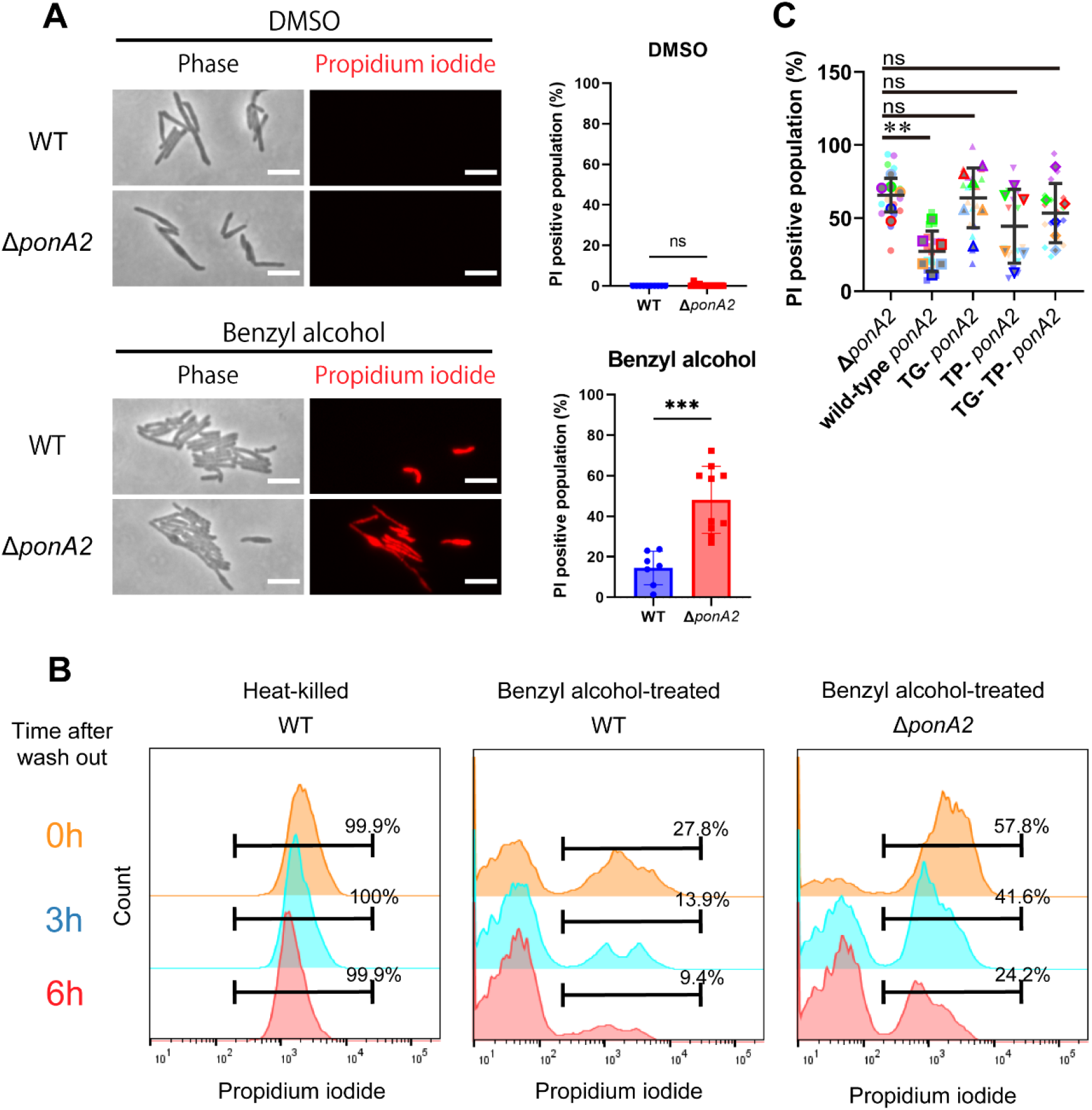
PonA2 promotes cell impermeability upon benzyl alcohol treatment. (A top) benzyl alcohol- or DMSO-treated wild-type or Δ*ponA2* strains were stained with propidium iodide to detect membrane-compromised cells. Scale bars, 5 µm. (A bottom) Populations of propidium iodide-positive cells in individual fields of wild-type (n = 160) or Δ*ponA2* (n = 327) are calculated and plotted. Mann-Whitney U-test. ***, p < 0.0005. (B) Flow cytometry analysis of propidium iodide-positive cells in time course after benzyl alcohol washout. Cells were incubated at 65℃ for one hour for heat-killed control. Data are representative of three independent experiments. (C) Δ*ponA2 M. smegmatis* complemented with wild-type *ponA2* or alleles with mutated transpeptidase (TP-) and/or transglycosylase (TG- and TP- TG-) domains were treated with benzyl alcohol and stained with propidium iodide. Propidium iodide staining quantitated in the same way as the right panel of (A). Data were combined from six individual experiments. Statistical significance was determined by the Kruskal–Wallis test, followed by Dunn’s multiple comparison test. **, p < 0.01. In total, 603 < n < 1355 cells were examined.

**Supplementary Table.**
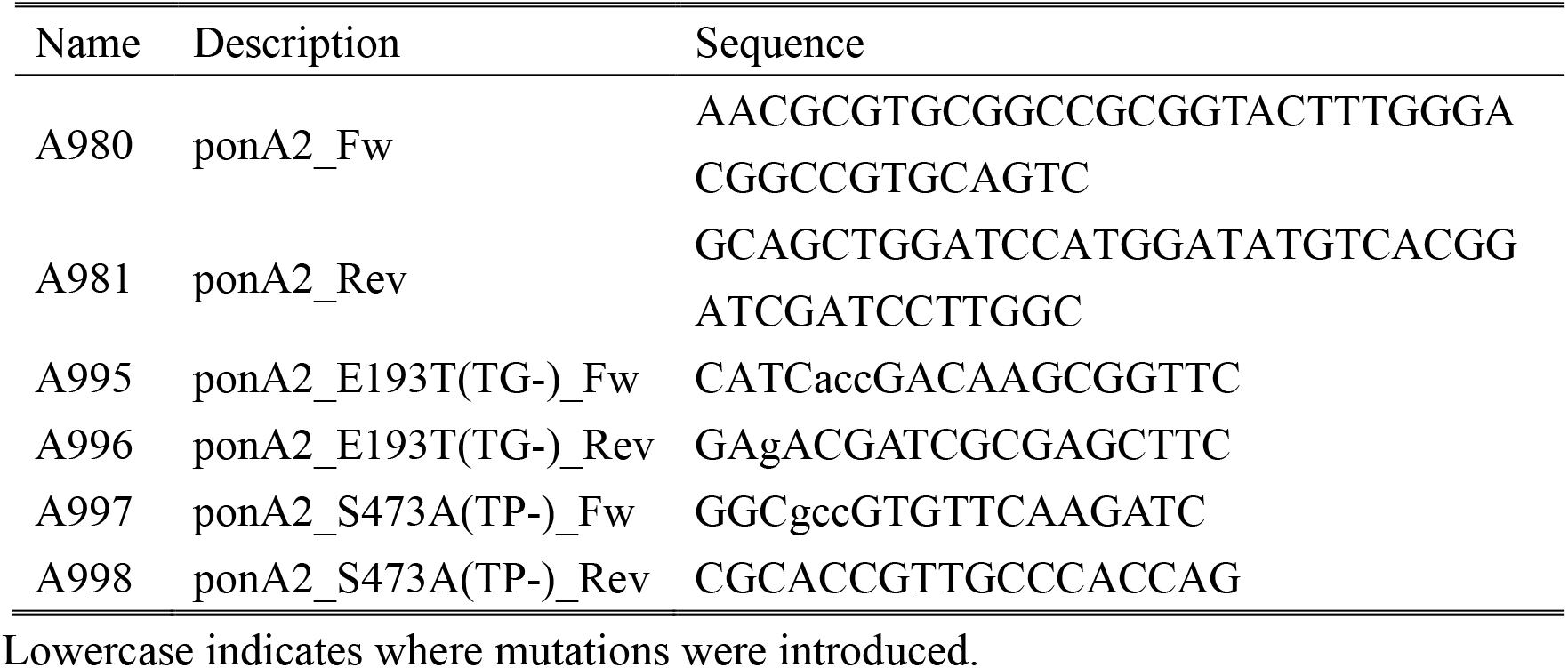
Primers used in this study.

## Notes

### Competing Interest Statement

The authors have declared no competing interest.

